# Mechanosensation Promotes Local Cell Wall Repair

**DOI:** 10.64898/2026.04.23.720310

**Authors:** Yannis Reignier, Dobrawa Amiri-Czajkowski, Remi Le Borgne, Nicolas Minc

## Abstract

Walled cells, such as those of fungi, plants and bacteria, must continuously survey the integrity of their protective cell wall (CW), as they grow, divide, or face mechanical challenges from their high internal turgor and environment. To date, however, how cells detect defects in their CW to activate local reinforcement or repair machineries remains unclear. We implemented a laser irradiation assay to locally wound the CW of model rod-shaped fission yeast cells. We found that laser irradiation causes a near instantaneous local thinning of the CW accompanied by the formation of a local bulge, a drop in turgor and growth arrest, followed by a progressive recovery of CW thickness over tens of minutes with cells eventually resuming growth. Remarkably, wounding the CW away from cell tips, caused the re-localization of canonical polarity regulators including the Rho GTPases Cdc42 and Rho1, that recruited actin assembly and vesicular trafficking of CW synthases to repair the CW. A candidate screen approach suggests that the transmembrane surface mechanosensors Wsc1 and Mlt2 detect local CW defects and promote the recruitment of the Rho GEFs Gef1 and Rgf1 that activate Cdc42 and Rho1. Therefore, these findings delineate a mechanochemical pathway, coupling surface mechanosensing to polarity regulation and CW synthesis, that defines how walled cells may detect and repair local defects in their CWs to safeguard surface integrity.

## INTRODUCTION

The Cell Wall is a thin elastic layer built from polysaccharides and proteins that encases the plasma membrane and protects the surface of plant, bacterial and fungal cells [1–4]. One of its primary function is to bear large mechanical stresses derived from substantial internal turgor pressure typical of walled cells [5–7]. As exemplified by the bursting phenotype of mutant cells defective in CW assembly, CW integrity is vital for walled cells [8–11]. However, the CW is dynamic in nature, changing shape, thickness and material properties to drive cell division, growth or cell-cell fusion, for instance [9,12–15]. To cope with such dynamic changes, walled cells have evolved surface sensing modules that may continuously monitor the mechanical state of the CW to adjust synthesis or remodeling [6,16–19]. Studies of these modules have led to the general proposal of mechano-transduction pathways that may couple surface sensing to downstream regulation of CW assembly. To date, however, how cells detect local mechanical changes in their CW to promote local reinforcement has remained elusive, in part because of a lack of quantitative assays to fragilize the CW in time and space.

Yeast and fungal cells target CW assembly and remodeling to specific cortical zones, to promote polarized growth and cell division [20]. This process is regulated by polarity modules that rely on the GTP-bound state of small GTPases such as Cdc42 or Rho1, regulated by GEFs and GAPs. In their active forms, these GTPases promote actin assembly and vectorial secretion of CW transmembrane remodeling enzymes. Fungal cells monitor the state of their CW through a conserved cascade called the Cell Integrity Pathway (CIP). This pathway has been well studied in the model yeasts *Saccharomyces cerevisiae* and *Schizosaccharomyces pombe* [17,21]. It encompasses surface mechanosensors of the Mid and Wsc families; single pass transmembrane proteins featuring extracellular domains that may directly interact with CW polysaccharides [10,22–24]. These domains are thought to directly sense modulations in CW thickness, stress or strain, through putative conformational changes, and trigger the downstream activation of the Rho GTPase Rho1 that promotes CW assembly, as well as a MAPK cascade, that turns on the transcription of CW remodeling genes. The rod-shape fission yeast, *S. pombe*, features two main surface sensors Wsc1 and Mtl2 that complement each other to support cell surface integrity and survival [10]. Previous studies have suggested that these sensors may interact through their cytoplasmic C-terminal with the Rho1 GEFs Rgf1 and Rgf2, to activate Rho1 and other downstream elements of the CIP [10]. Yet, it remains unclear how these sensors may detect local CW mechanical defects to influence polarity modules and spatially regulate CW repair or reinforcement.

Here, inspired by previous assays in *S. cerevisiae* and *S. pombe* [25–27], we implemented a robust assay based on UV laser irradiation to create controlled damages around the CW of fission yeast cells. By titrating laser power to create subtle damages that still permit cell survival, we monitored CW reparation mechanisms. We found that laser irradiation causes the CW to locally thin near instantaneously, yielding to the formation of a bulge, a drop in turgor and a global cell growth arrest. By monitoring polarity and CIP proteins with live imaging following CW wounding, we found that most of these factors are rerouted to the wound site within tens of minutes to promote CW repair. Accordingly, the CW recovered its initial thickness over similar time frames with cells resuming growth at cell tips. By performing a candidate screen, we demonstrate that CW damage may be sensed as a mechanical defect by both surface sensors Wsc1 and Mtl2, that recruit the GEFs Rgf1 and Gef1 to promote the activation of Rho1 and Cdc42 and drive local actin assembly and vesicular trafficking of CW remodeling enzymes that repair the CW. Therefore, this study identifies dynamic hierarchical mechanisms that couple surface mechanosensation in the CW to cell polarity to promote local CW repair and ensure cell surface integrity.

## RESULTS

### A laser-based assay to locally damage the CW

We sought to design an assay to study the ability of walled cells to locally repair CW damages. Using the rod-shaped fission yeast as a model, we implemented a local laser irradiation assay based on focusing a UV laser spot of ∼2.5 µm^2^ on the cell surface [27]. By systematically tuning laser power, we identified the maximum irradiation level that did not cause cell death but that significantly affected the CW locally. The impact of laser irradiation on the CW was readily visible in the formation of a local bulge when the laser was targeted to cell sides. This bulge formed within few tens of seconds in 100% of irradiated cells and was irreversible (Fig 1A-D; and Fig S1A-S1B). To assess the state of the CW, we wounded cells with the laser beam and immediately fixed them to perform Correlative Light and Electron Microscopy (CLEM) and image the CW at high resolution. The bulge was clearly visible in TEM images and was marked by a significant decrease of ∼40% in CW thickness as compared to the opposite untouched cell side, reflecting a significant local alteration of the CW (Fig 1E-F and Fig S1C). We also documented the kinetic of CW thickness changes following laser irradiation, using a sub-resolution fluorescence-based method to measure CW thickness in live cells [9] (Fig 1G). In agreement with CLEM results, this analysis showed that the CW was thinner at the wound site as compared to the rest of the cell, supporting that local laser irradiation can be used to locally thin and damage the CW in living cells. Remarkably, this live assay also revealed a progressive recovery of CW thickness over tens of minutes with a complete recovery achieved within 60 to 80 minutes, indicating the existence of a local CW repair mechanism. (Fig 1G-H, Fig S1D and Movie S1).

**Figure 1.**
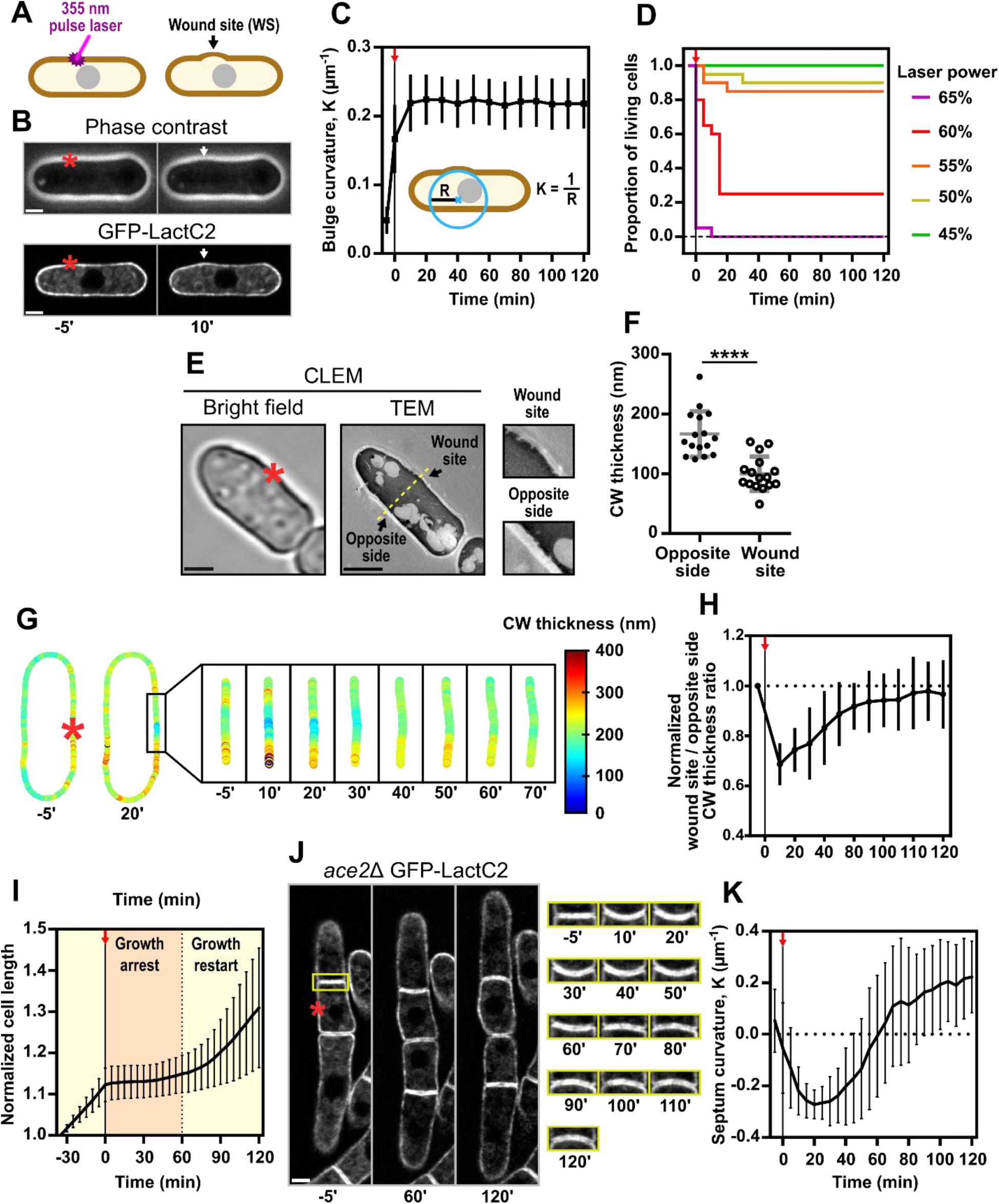
A Laser based assay to locally damage the Cell Wall. **(A)** Scheme illustrating the cell wall wounding assay used throughout the study. **(B)** Top panel: Phase contrast image of a yeast cell before and after laser irradiation. The laser irradiation site is marked by a red asterisk. Bottom panel: Midplane confocal image of the same cell expressing the plasma membrane marker GFP-LactC2. White arrows point to the bulge emerging after laser irradiation. **(C)** Kinetic of plasma membrane curvature at the wound site (WS): time zero was acquired few seconds after laser irradiation and is marked by a red arrow (n = 18 cells). **(D)** Cell survival after laser irradiation for different laser power (n=20 cells for each condition). **(E)** Corelative Light and Electron Microscopy (CLEM) of an ablated cell. Left panel: bright field image of a cell just after laser irradiation. Right panel: Transmission Electron Microscopy (TEM) image of the same cell. The yellow dotted line illustrates the axis used to measure cell wall (CW) thickness in panel G. **(F)** CW thickness measurements at the WS and the opposite side performed on CLEM-TEM images (n=16 cells). **(G)** CW thickness map of a living cell before and after laser irradiation computed from confocal images of a GFP-LactC2 expressing cell labeled with lectin-Alexafluor647 (see methods). **(H)** Kinetic of CW thickness at the WS normalized to the opposite side extracted from CW thickness maps (n=24 cells). **(I)** Cell growth before and after laser irradiation (n=30 cells). **(J)** Left panel: confocal timelapse of *ace2*Δ cells expressing GFP-LactC2 before and after laser irradiation. Right panel: timelapse of the septum between two of those cells. **(K)** Kinetic of septum curvature in *ace2*Δ cells expressing GFP-LactC2 before and after laser irradiation. (n=10). All error bars correspond to standard deviations (SD). Results were compared using a Wilcoxon matched-pairs signed rank test (****; P value < 0.0001). Scale bars: 2 µm.

We next monitored the more global impact of CW wounding caused by laser irradiation. We first performed dynamic cell length measurements to quantify polar cell growth in long time-lapses. Interestingly, this showed that cells completely halt growth within ∼5 minutes after laser irradiation, but that they eventually resume growth after ∼60-80 minutes, in concomitance with CW thickness recovery at the wound (Fig 1H-I and Fig S1B). Next, we assayed the turgid state of wounded cells. For this, we wounded septating cells, expressing the membrane marker GFP-LactC2, and observed that the septum always bend towards the wounded cell, suggesting a drop of turgor in the wounded cell [28,29] (Fig S1E). To monitor turgor recovery over longer time periods, we used an *ace2*Δ mutant defective in cell separation upon septation [30] (Fig 1J). This showed that the septum was initially flat and started to bend toward the wounded cell few minutes following laser irradiation, reaching a maximum curvature at around ∼20 min, after which septum curvature decreased to go back to 0 at around ∼60 min, before eventually bending toward the opposite cell compartment (Fig 1J-K). This kinetic of septum bending confirms a rapid turgor drop following CW wounding, and suggests a progressive recovery concomitant with CW thickness recovery, that eventually lead to an overshoot in turgor values in the wounded cell. Importantly, the initial drop in turgor pressure suggested leaks across the membrane at the wounded site, which we confirmed using propidium iodine staining [31] (Fig S1F-G). Altogether, these data indicate that local CW wounding causes an initial growth arrest and turgor drop, followed by a progressive recovery over ∼1h concomitant and potentially linked to a local repair of the CW.

### Cell wall damage triggers a repair mechanism

Fission yeast cells are highly polarized with canonical polarity and CW regulating proteins localized to growing cell tips during interphase and to the cell middle during septation [20]. Given the observed recovery of CW thickness following wounding on cell sides, we asked if polarity factors normally located to cell tips would be redirected to the wound site. Using live spinning disk confocal microscopy, we first monitored the localization of GFP-Bgs1 and GFP-Bgs4, two β-glucan synthase catalytic subunits, as well as that of the α-glucan synthase Ags1-GFP, all crucial for CW synthesis [32–34]. Strikingly, CW wounding by laser irradiation caused these factors to progressively detach from cell tips and to reaccumulate at the wound site over the course of ∼30 min, forming an ectopic polar domain of similar size as that found at cell tips in normal cells. Interestingly, however, this accumulation was transient, as this ectopic domain eventually disassemble to re-localize to incipient growing tips after ∼60 min presumably promoting the restart of tip growth as documented above (Fig 2B-E, Fig S2A-D and Movie S2). Importantly, targeting the laser beam to cell tips, also caused a transient detachment of the glucan synthase polar domain, followed by a more rapid reformation at the same tip (Fig S2E-G). Finally, we also noted that glucan synthase accumulation to the wound site was significantly delayed in septating cells, suggesting a differential competition between wound recruitment with cell tips vs septa (Fig S2H-J). Together, these findings suggest that cells actively detect local wounds in their CW to promote the redirection of key CW enzymes that may re-synthetize CW material to repair the CW.

**Figure 2.**
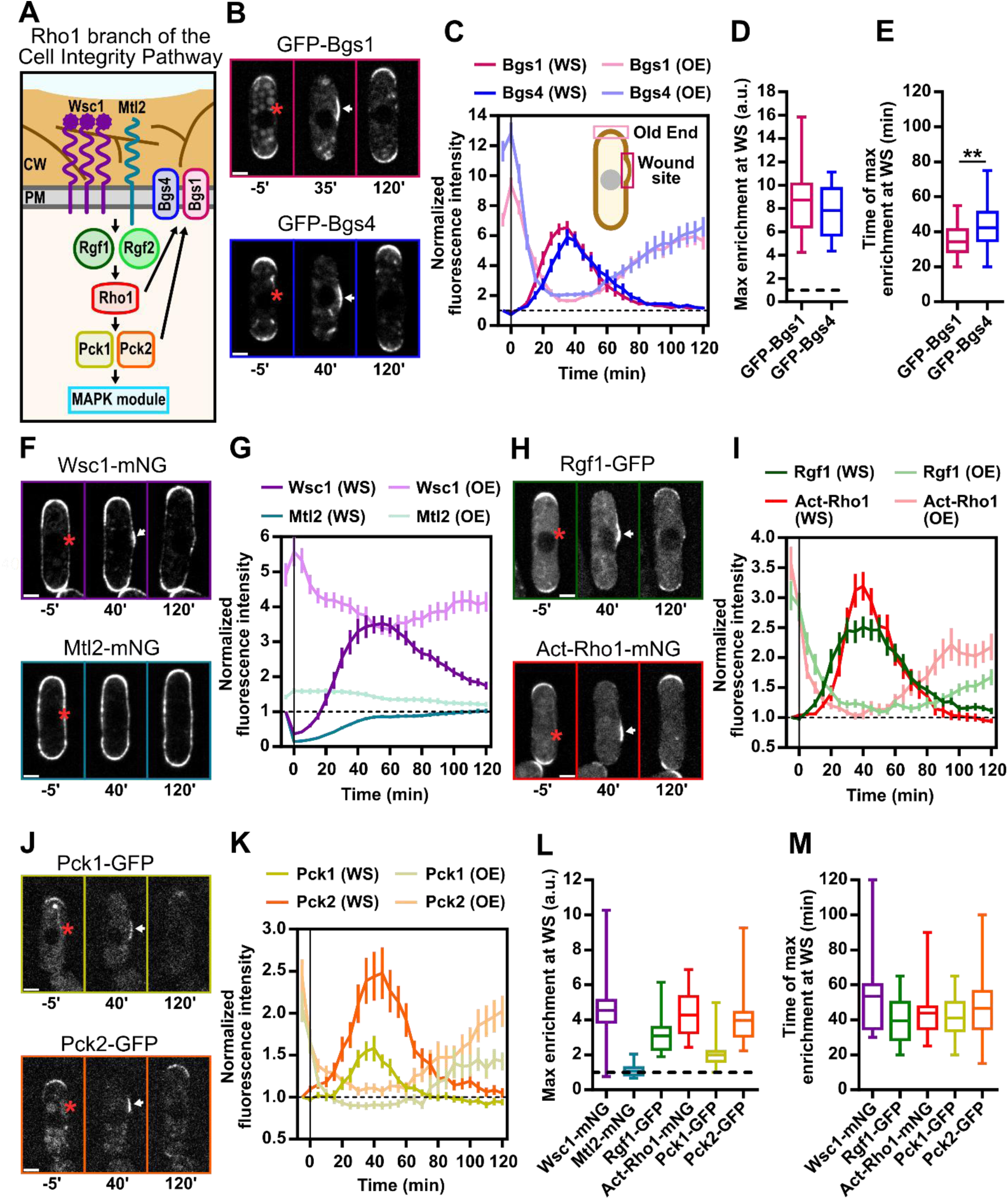
Cell wall damage triggers local activation of the Cell Integrity Pathway. **(A)** Scheme of the Rho1 branch of the Cell Integrity Pathway (CIP). **(B)** Confocal timelapses of cells expressing: GFP-Bgs1 (top panel) or GFP-Bgs4 (bottom panel), before and after laser irradiation. Laser irradiation sites are marked by red asterisks. White arrows point to protein accumulation zones at the wound site (WS). **(C)** Kinetics of GFP-Bgs1 and GFP-Bgs4 signal intensity at the WS and the old end (OE): time zero was acquired few seconds after laser irradiation. (n=30 cells for each strain). **(D)** Mean maximum enrichment of GFP-Bgs1 and GFP-Bgs4 measured at the WS. A value of 1 corresponds to the initial fluorescence signal measured at the WS before laser irradiation**. (E)** Mean timing of maximum GFP-Bgs1 and GFP-Bgs4 enrichment at the WS. **(F)** Confocal timelapses of cells expressing: Wsc1-mNG (top panel) or Mtl2-mNG (bottom panel), before and after laser irradiation. **(G)** Kinetics of Wsc1-mNG and Mtl2-mNG signal intensity at the WS and the OE (n=30 cells for each strain). **(H)** Confocal timelapses of cells expressing: Rgf1-GFP (top panel) or Active-Rho1-mNG (RBD-mNG, bottom panel), before and after laser irradiation. (I) Kinetics of Rgf1-GFP and Active-Rho1-mNG signal intensity at the WS and the OE (n=30 cells for each strain). **(J)** Confocal timelapses of cells expressing Pck1-GFP (top panel) or Pck2-GFP (bottom panel), before and after laser irradiation. **(K)** Kinetics of Pck1-GFP and Pck2-GFP signal intensity at the WS and the OE. **(L)** Mean maximum enrichment of the Cell Integrity Pathway (CIP) proteins measured at the WS. **(M)** Mean timing of maximum enrichment for the protein of the CIP at the WS. Kinetics error bars show standard error of mean (SEM). Whiskers show full data range. Results were compared using the two-tailed Mann-Whitney test (**, P value < 0.01). Scale bars: 2 µm.

### Cell wall damage triggers the local activation of the Cell Integrity Pathway

Glucan synthases may be activated by the CIP, raising the possibility that components of the CIP may also be retargeted to CW wound sites [25] (Fig 2A and Movie S2). We first monitored Wsc1 and Mtl2, as these CW sensors have been described as upstream activators of the Rho1 branch of the CIP [10]. Mtl2-mNG was already present all around the cell contour and did not become enriched at the wound site, beyond a slight recovery following laser irradiation-induced photo-bleaching. In contrast, Wsc1-mNG partially depolarized from cell tips to reaccumulate at the wound site ∼30-40 min following laser irradiation (Fig 2F-G, 2L-M and Movie S2). In addition, Wsc1-mNG appeared to form small clusters decorating the wound site, reminiscent of those formed upon CW compression, suggesting that this sensor may be locally active [24]. We conclude that both Wsc1 and Mtl2 mechanosensors may be already present or actively recruited at CW damage sites, to potentially activate downstream CIP effectors.

To confirm the local activation of the CIP, we conducted observations on downstream effectors (Fig 2A). We first monitored the localization of Rgf1-GFP and Act-Rho1-mNG, a construct that specifically recognizes the active GTP-bound form of Rho1 [35]. Both were redirected to the wound site with a similar kinetic as glucan synthases described above (Fig 2H-I; Fig 2M and Movie S2). As active Rho1 serves as a regulatory subunit for glucan synthases, this observation suggests that the whole complexes promoting CW synthesis may be present at the wound site to repair the CW. Finally, the downstream CIP effectors Pck1-GFP and Pck2-GFP, also followed a similar re-localization dynamic to the wound site (Fig 2J-M). Therefore, many core components of the CIP may be rerouted to the wound site, indicating an active local mechanosensation of CW damage and local activation of a repair pathway.

### Cdc42-based polarity and F-actin assembly are rerouted to the wound site

Glucan synthases are transmembrane proteins that are transported to the cell surface within secretory vesicles trafficking along actin cables [33,34,36]. We thus aimed to test if actin-regulators and other polarity components could also be rerouted to the wound site (Fig 3A). We first imaged the localization of the active GTP-bound form of Cdc42 a major regulator of cell polarity and actin assembly in fission yeast, using the biosensor CRIB-3GFP [37]. Alike CIP components and glucan synthases, Cdc42 activity was redirected to the wound site after laser irradiation with a maximum activation at around 35 minutes, followed by a recovery at cell tips. The formin For3, a downstream effector of Cdc42 that promotes actin cable polymerization, was also recruited to the wound site after laser irradiation (Fig 3B-C and 3H-I). Accordingly, directly imaging F-actin with the LifeAct-mGFP probe revealed a partial depolarization from the cell tips and the formation of ectopic actin patches and cables growing from the CW wound site. These cables promoted trafficking towards the wound site as demonstrated by the ectopic accumulation of the myosin type-V motor Myo52 (Fig 3D-E and 3H-I). These findings demonstrate that Cdc42-based polarized actin assembly is also rerouted towards the wound site to promote vectorial secretion of transmembrane enzymes there.

**Figure 3.**
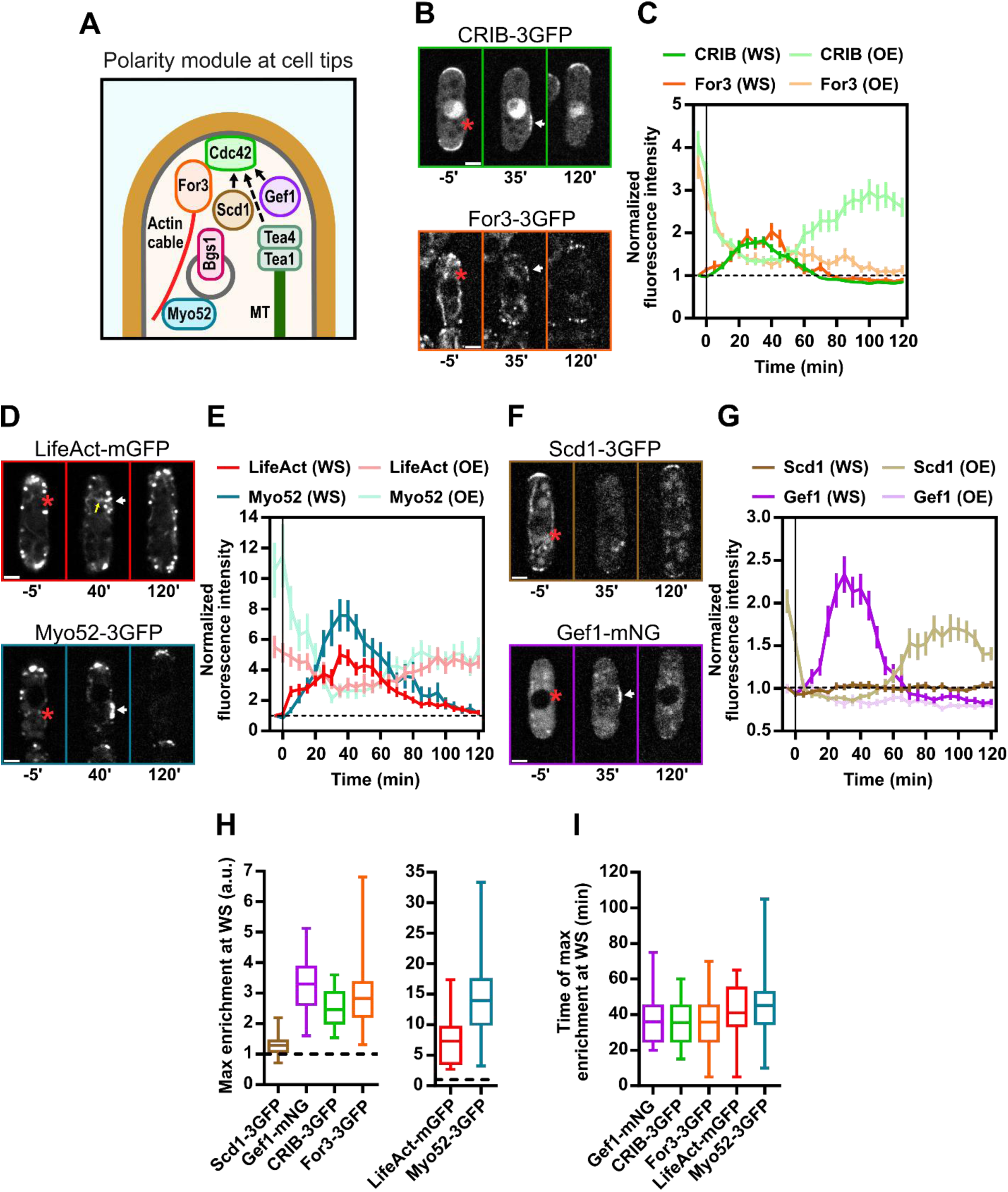
The polarity machinery is rerouted to the wound site. **(A)** Schematic representation of cell polarity regulators at cell tips. **(B)** Confocal timelapses of WT cells expressing CRIB-3GFP (top panel) or For3-3GFP (bottom panel) before and after laser irradiation. Laser irradiation sites are marked by red asterisks. White arrows point to protein accumulation zones at the wound site (WS). **(C)** Kinetics of CRIB-3GFP and For3-3GFP signal intensity at the WS and the old end (OE): time zero was acquired few seconds after laser irradiation (n=30 cells for each strain). **(D)** Confocal timelapses of WT cells expressing LifeAct-mGFP (Top panel) or Myo52-3GFP (Bottom Panel) before and after laser irradiation. **(E)** Kinetics of LifeAct-mGFP and Myo52-3GFP signal intensity at the WS and the OE. (n=30 cells in each strain). **(F)** Confocal timelapses of WT cells expressing Scd1-3GFP (Top Panel) or Gef1-mNG (Bottom Panel), before and after laser irradiation. **(G)** Kinetics of Rgf1-GFP and Active-Rho1-mNG signal intensity at the WS and the OE (n=30 cells in each strain). (H) Mean maximum enrichment of polarity factors measured at the WS. The value of 1 corresponds to the initial fluorescence signal measured at the WS before laser irradiation. (For each condition: n=30 cells). (I) Mean timing of maximum enrichment of polarity factors at the WS. (For each condition: n=30). Kinetics error bars show standard error of mean (SEM). Whiskers show full data range. Scale bars: 2 µm.

In order to understand how Cdc42 may be activated at the wound site, we next imaged Scd1 and Gef1, two Rho GEFs known to activate Cdc42 in *S. pombe* [38]. Remarkably, Scd1-3GFP which normally exhibit a marked polarized distribution to cell tips, detached from cell tips following laser irradiation, but did not exhibit any accumulation to the wound site, suggesting it is dispensable to activate Cdc42 there. In sharp contrast, Gef1-mNG was mostly cytoplasmic before laser irradiation, with slight accumulation at cell tips, but became highly enriched at the wound site following laser irradiation, to eventually vanish in the cytoplasm following CW repair (Fig 3F-G: Movie S3). Those results align with the notion that Gef1 may be the main GEF promoting Cdc42 activation on cell sides [39,40], and suggest that Gef1 may locally activate Cdc42 at the wound site to reroute actin-based trafficking and promote the local secretion of transmembrane enzymes like glucan synthase that drive CW repair.

### F-actin, but not microtubules, is required to localize glucan synthases to the wound site

To begin to dissect mechanisms of localized CW repair at the wound site, we first sought to test the role of the cytoskeleton. Unlike F-actin, the microtubule (MT) cytoskeleton did not seem to reorganize following CW wounding by laser irradiation. Accordingly, polarity components that depend on MT-based transport, like Tea1 did not exhibit any change in localization following laser irradiation (Fig S3A-B). In addition, depolymerizing MTs with low doses of Methyl Benzimidazole-2-yl Carbamate (MBC) did not affect the dynamics of recruitment of GFP-Bgs4 to CW wound sites. Similar results were also obtained in a *tea1*Δ mutant, although we noted significant defects in the reformation of the polar GFP-bgs4 domain at cell tips upon CW repair in this mutant (Fig 4A-B and Fig S3C-E). These results suggest that MT-based polarity is dispensable for targeting CW assembly to ectopic sites in this context [41,42].

**Figure 4.**
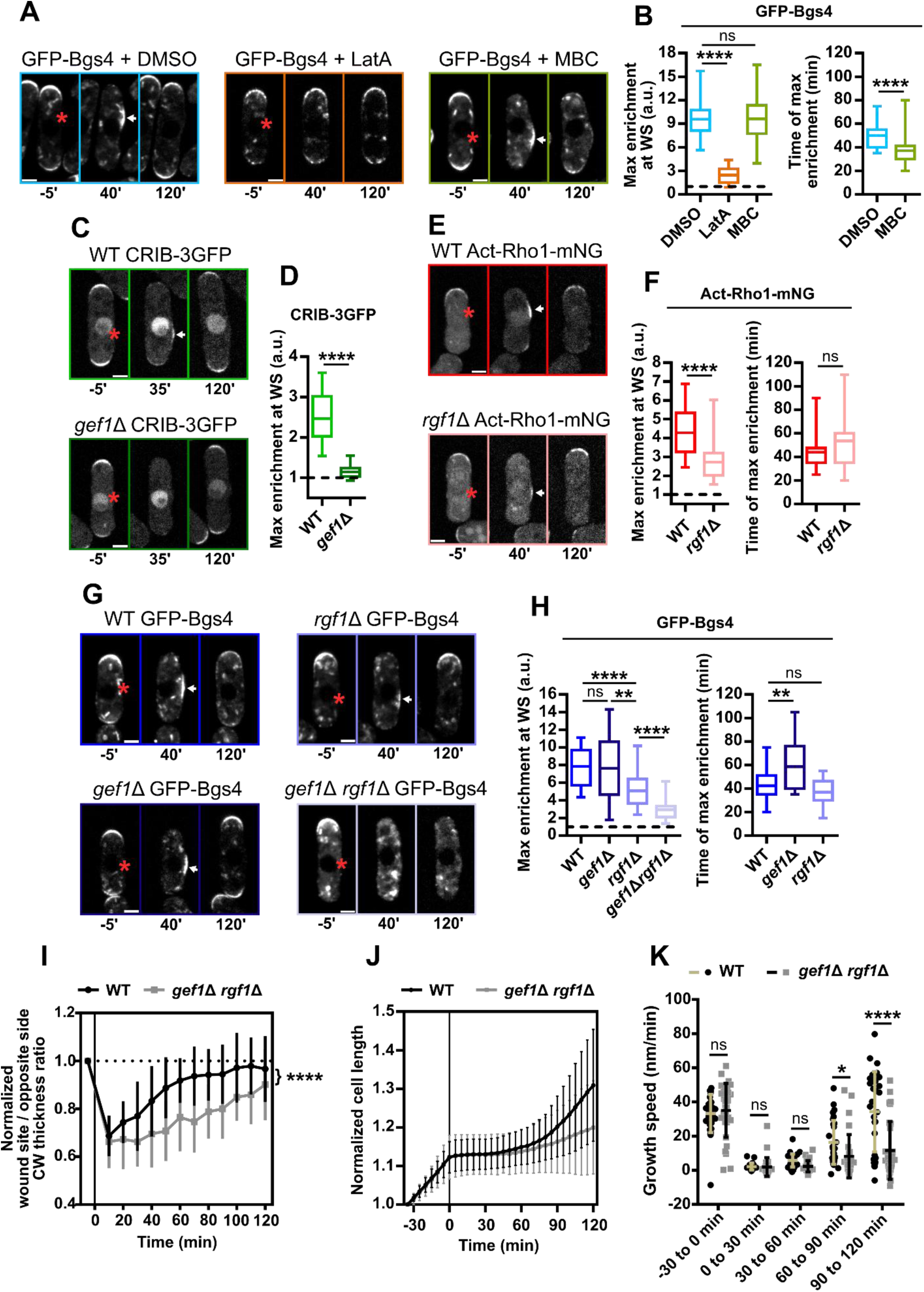
Rho1 and Cdc42 recruit glucan synthases to the wound in an actin dependent manner. **(A)** Confocal timelapses of cells expressing GFP-Bgs4 treated with: DMSO; 100 µM Latrunculin A; or 25 µg/ml MBC, before and after laser irradiation. Laser irradiation sites are marked by red asterisks. White arrows point to protein accumulation zones at the wound site (WS). **(B)** Left panel: mean maximum enrichment of GFP-Bgs4 measured at the WS. (n=30, 20 and 30 cells respectively). The value of 1 corresponds to the initial fluorescence signal measured at the WS before laser irradiation. Right panel: mean timing of maximum GFP-Bgs4 enrichment at the WS. **(C)** Confocal timelapses of a WT cell (Top panel) or a *gef1*Δ mutant cell (Bottom panel), expressing CRIB-3GFP before and after laser irradiation. **(D)** Left panel: mean maximum enrichment of CRIB-3GFP measured at the WS. Right panel: mean timing of maximum CRIB-3GFP enrichment at the WS. **(E)** Confocal timelapses of a WT cell (top panel) or a *rgf1*Δ mutant cell (bottom panel), expressing Active-Rho1-mNG before and after laser irradiation. **(F)** Left panel: mean maximum enrichment of Active-Rho1-mNG measured at the WS. (n=30 and 40 cells respectively). Right panel: mean timing of maximum Active-Rho1-mNG enrichment at the WS. **(G)** Confocal timelapses of a WT cell (top left panel), a *gef1*Δ mutant cell (bottom left panel) a *rgf1*Δ mutant cell (top right panel), and a *gef1*Δ *rgf1*Δ double mutant cell (bottom right panel), all expressing GFP-Bgs4 before and after laser irradiation. **(H)** Left panel: mean maximum enrichment of GFP-Bgs4 measured at the WS. (n=30 cells for all strains). Right panel: mean timing of maximum GFP-Bgs4 enrichment at the WS. **(I)** CW thickness kinetics at the WS for WT cells (in black) and *gef1*Δ *rgf1*Δ double mutant cells (in grey) (n=24 and 15 cells respectively). **(J)** Cell growth of WT cells (in black) and *gef1*Δ *rgf1*Δ double mutant cells (in grey), before and after laser irradiation, marked by the vertical line a time=0 min (n=30 cells for both strains). **(K)** Growth elongation speed of WT cells (in black) and *gef1*Δ *rgf1*Δ double mutant cells (in grey), binned on different time periods. Whiskers show full data range. Kinetics and growth rate error bars correspond to SD. Results in B, D, F and H were compared using the Mann-Whitney test. Results in I and K were compared using the two-way ANOVA test (*p < 0.05; **p < 0.01; ****p < 0.0001; ns: not significant). Scale bars: 2 µm.

In sharp contrast, depolymerizing F-actin with low doses of Latrunculin A (Lat A) completely blocked the re-localization of GFP-Bgs4 to CW wounds on cell sides, with cells retaining polar domains at cell tips throughout the process (Fig 4A-B). As a consequence of this lack of retargeting CW synthases to the wound site, cells treated with Lat A died progressively after laser irradiation, indicating that proper F-actin mediated CW synthase rerouting may be key to support cell surface integrity (Fig S3F).

### The Rho GEFs Gef1 and Rgf1 complement each other to regulate local CW repair

The active forms of Rho-GTPases Cdc42 and Rho1 are important regulators of F-actin assembly. Therefore, in order to investigate how F-actin is regulated to promote glucan synthases polarization to the wound site, we monitored the localization of active Cdc42 and Rho1 in mutants lacking some of their GEFs. In agreement with the recruitment of Gef1, but not Scd1, to the wound site, we found that *gef1*Δ cells could not redirect CRIB-3GFP to the wound site after laser irradiation (Fig 4C-D). In contrast, a *rgf1*Δ mutant lacking the major Rho1GEF, showed only a slight decrease in Cdc42 activity at the wound site (Fig S4A-B). Interestingly, both *rgf1*Δ and *gef1*Δ exhibited a marked reduction in the level of active-Rho1 recruitment to the wound site, suggesting a plausible regulation of Rho1 activity from Cdc42-based activity in this context (Fig 4E-F; Fig S4C-D). However, the *rgf1*Δ mutant presented a stronger reduction in Rho1 activation, similar to the level of a double *gef1*Δ *rgf1*Δ mutant, indicating that Rgf1 is still the main regulator of Rho1 activity in this process (Fig S4 C-D).

These results prompted us to test the role of these GEFs on the recruitment of glucan synthases. Strikingly, a *gef1*Δ mutant, but not a *rgf1*Δ, exhibited a strong reduction in GFP-Bgs1 enrichment to the wound site while conversely a *rgf1*Δ mutant, but not *gef1*Δ affected GFP-Bgs4 enrichment (Fig 4G-H; Fig S4E-F; Movie S4). This differential regulation of Bgs1 and Bgs4 is reminiscent of divergent mechanisms of glucan synthase recruitment at the septum [43,44], and indicate that both Rho GTPases cooperate to ensure the proper recruitment of glucan synthases. Accordingly, *gef1*Δ mutants, although defective in activating Cdc42 at the wound site, still exhibited a rerouting of actin assembly, suggesting a potential additional regulation of F-actin polarization by Rho1 or other GTPases [45,46] (Fig S4G-H). Furthermore, glucan synthase recruitment was nearly abolished in a double *gef1*Δ *rgf1*Δ mutant, leading to major delays in CW rethickening and polar growth resumption after CW wounding (Fig 4G-K and Movie S4). Overall, these results suggest that the active forms of both Cdc42 and Rho1 promote the recruitment of the Bgs1 and Bgs4 glucan synthases to the wound site, with Cdc42 preferentially influencing Bgs1 and Rho1influencing Bgs4.

### CW damages are detected as mechanical defects by surface mechanosensors

The CW is put under large tensional stress by turgor pressure [4,28]. This stress is inversely proportional to CW thickness, so that the thinning created by the laser could in principle result in local enhanced stress that may be detected by CW surface sensors (Fig 5A). To test this hypothesis, we used a *gpd1*Δ mutant deficient in turgor pressure adaptation and rinsed cells with media supplemented with 0.5 M sorbitol, a dose that was previously shown to abolish turgor [27,47,48]. We irradiated these deflated cells and compared the recruitment of Wsc1-mNG to control cells placed in normal isotonic medium. Strikingly, while untreated *gpd1*Δ cells exhibited a re-localization of Wsc1-mNG to the wound site similar to WT cells, deflated cells in sorbitol failed to re-localize Wsc1-mNG to the site of laser irradiation. A similar lack of recruitment in deflated cells was also observed for the sensors of GTP-bound Cdc42 and Rho1, CRIB-3GFP and Act-Rho1, as well as for the glucan synthase GFP-Bgs4 (Fig 5B-D). These results indicate that CW damages may be detected as a local mechanical stress enhancement that could promote the accumulation of surface CW sensors to activate and recruit downstream polarity and CW regulator effectors [24].

**Figure 5.**
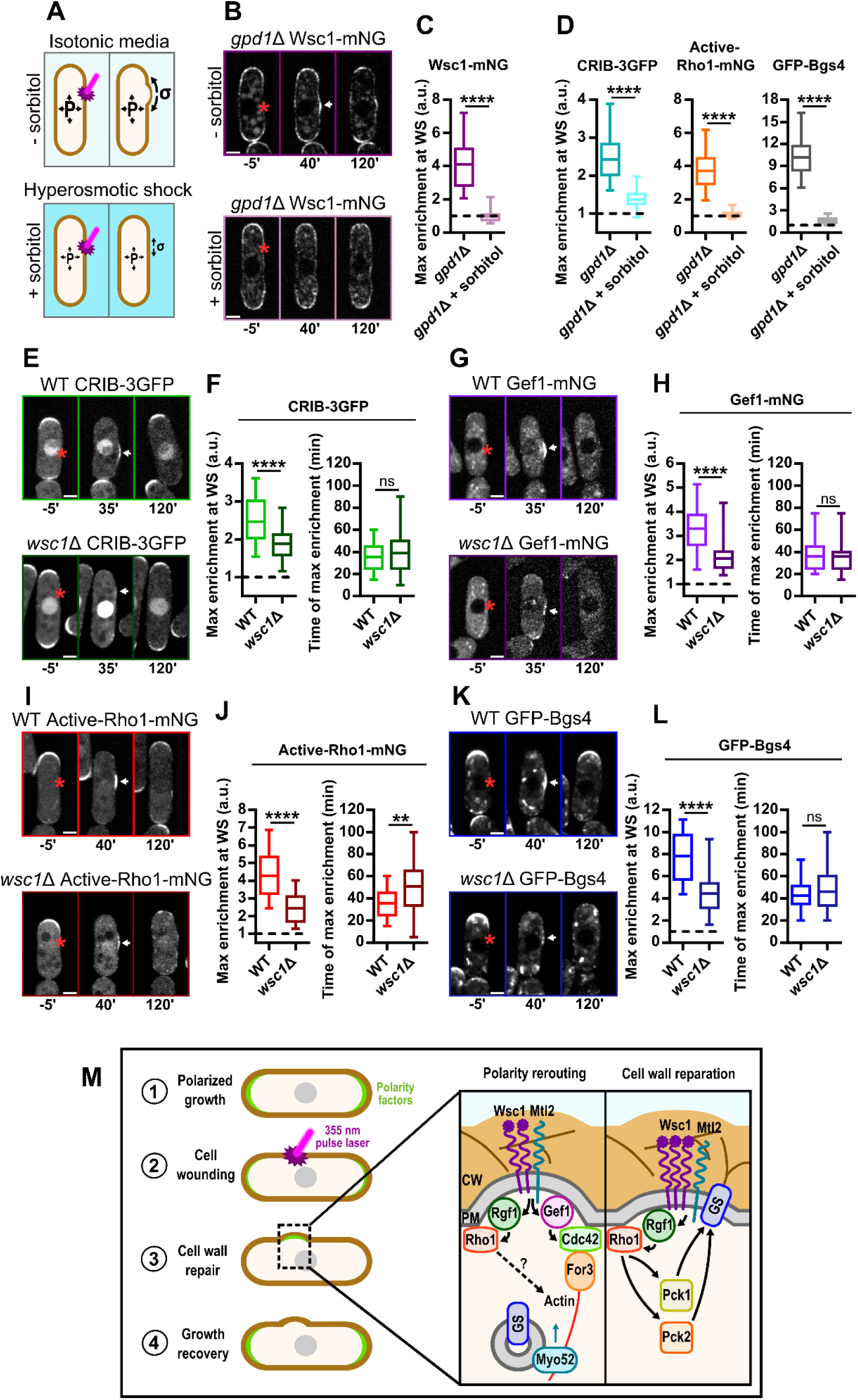
The Wsc1 mechanosensor detects local CW stress to reroute cell polarity to the wound site. **(A)** Scheme illustrating the impact of turgor pressure on the local lateral stress at the damaged CW. Dark blue symbolizes hyperosmotic media. Decrease of turgor pressure P leads to a diminished local CW stress σ after local laser damage. **(B)** Confocal timelapses of an untreated *gpd1*Δ mutant cell (top panel) or a *gpd1*Δ mutant cell incubated in 0.5 M sorbitol (bottom panel) expressing Wsc1-mNG before and after laser irradiation. Laser irradiation sites are marked by red asterisks. The white arrows point to protein accumulation zones at the wound site (WS). **(C)** Mean maximum enrichment of Wsc1-mNG measured at the WS (n=30 cells in both conditions). The value of 1 corresponds to the initial fluorescence signal measured at the WS before laser irradiation. **(D)** Left panel: mean maximum enrichment of CRIB-3GFP measured at the WS; middle panel: mean maximum enrichment of Active-Rho1-mNG measured at the WS; right panel: mean maximum enrichment of GFP-Bgs4 measured at the WS; in absence or presence of 0.5 M sorbitol (n=30 cells for all strains and conditions). **(E)** Confocal timelapses of a WT cell (top panel) or a *wsc1*Δ mutant cell (bottom panel), expressing CRIB-3GFP. **(F)** Left panel: mean maximum enrichment of CRIB-3GFP measured at the WS. Right panel: mean timing of maximum CRIB-3GFP enrichment at the WS. (n= 30 and 33 cells respectively). **(G)** Confocal timelapses of a WT (top panel) or a *wsc1*Δ mutant cell (bottom panel) expressing Gef1-mNG. (H) Left panel: mean maximum enrichment of Gef1-mNG measured at the WS. Right panel: mean timing of maximum Gef1-mNG enrichment at the WS (n= 30 cells for each strain). **(I)** Confocal timelapses of a WT cell (top panel) or a *wsc1*Δ mutant cell (bottom panel) expressing ActiveRho1-mNG. (J) Left panel: mean maximum enrichment of ActiveRho1-mNG measured at the WS. Right panel: mean timing of maximum ActiveRho1-mNG enrichment at the WS. (n= 30, cells in both strains). **(K)** Confocal timelapses of: a WT cell (top panel) or a *wsc1*Δ mutant cell (bottom panel) expressing GFP-Bgs4. **(L)** Left panel: mean maximum enrichment of GFP-Bgs4 measured at the WS. Right panel: mean timing of maximum GFP-Bgs4 enrichment at the WS. (n= 30 cells for each strain). **(M)** Proposed model of a mechanotransduction pathway for local CW repair, coupling CW mechanosensation to cell polarity and CW synthesis regulators. Whiskers show full data range. Results were compared using the Mann-Whitney test (**p < 0.01; ****p < 0.0001, ns: not significant). Scale bars: 2 µm.

To more directly test this hypothesis, we monitored the recruitment of several downstream effectors of polarity and CW synthesis in a *wsc1*Δ mutant. Strikingly, CRIB-3GFP recruitment to the wound site was significantly reduced in *wsc1*Δ, and a similar effect was also obtained for Gef1-mNG, indicating that sensor-mediated recruitment of Cdc42 activity may be mediated by Gef1. Furthermore, Wsc1 also largely influenced the recruitment of active-Rho1 and GFP-Bgs4 to the damage site (Fig 5E-L and Movie S5 and S6). We suspected that Mtl2 could be responsible for the residual recruitments observed in *wsc1*Δ. To test this, we monitored the recruitment of the same factors in the *mtl2*Δ single mutant, as well as in an inducible *wsc1*Δ *nmt1-mtl2* shut-off strain to deplete both sensors from cells [10]. The *mtl2*Δ mutant only sowed mild defects, while the recruitment of downstream factors was nearly abolished in the double shut-off strain (Fig S5A-H; Movie S5 and S6). This indicates that while this response is dominated by Wsc1, Mtl2 acts redundantly and can partially compensate for the loss of Wsc1. Finally, we tested the impact of affecting the CIP at a downstream level, by monitoring GFP-bgs4 recruitment in *pck2*Δ and *pmk1*Δ mutants. These mutants exhibited a significant reduction in glucan synthase recruitment to the wound, that was nonetheless higher than in *wsc1*Δ, suggesting that downstream transcriptional regulation of the CIP pathway may contribute, but that most of the recruitment of downstream polarity and CW regulators is directly mediated by surface sensors (Fig S5I-J). All together, these findings support a mechanosensing mechanism in which the surface sensors Wsc1 and Mtl2 may directly sense CW damages as mechanical signals, that they relay by recruiting the GEFs Gef1 and Rgf1 to prime local reassembly of polarity, F-actin and CW regulators needed for CW repair (Fig 5M).

## DISCUSSION

### Cell Wall dynamics and mechanosensing

Once considered as an inert barrier, the CW is being more and more recognized as a dynamic structure that can rapidly change material properties to promote surface integrity in physiological instances such as growth, division, reproduction or infection [13,49–51]. These adaptations often rely on surface mechanosensing that serve to detect particular mechanical challenges on the CW [6,7]. These challenges may emerge from intracellular reorganization such as during polar growth or septation, or from mechanical constraints in the environment. For instance, in both budding and fission yeasts, CW remodeling needed to drive polar mitotic growth or cell-cell fusion during mating, implicate mechanosensors of the Wsc or Mid family as well as downstream CIP activation [9,11,52–54]. These sensors also prime CW integrity in response to chemical, mechanical or osmotic perturbations in the environment [19,24,55,56]. Surface mechanosensing is also of fundamental importance for higher eukaryotic walled cells, such as those of plants: Growing pollen tubes use the surface mechanosensor BUPS1 to promote CW stiffening and survival, as they navigate through narrow gaps within stylar transmitting female tissues [57], and root hairs exploit the mechanoreceptor-like kinase FERONIA to promote CW integrity when growing within hard substrates such as soils [58]. Here, using a quantitative laser irradiation assay to fragilize the CW in time and space, we specifically monitored mechanisms of CW repair. We found that cells may detect local wounds through the surface mechanosensors, Wsc1 and Mtl2, that recruit downstream effectors of cell polarity and CIP to repair the CW. Therefore, our study adds to a global appreciation of the central role of mechanosensing for CW dynamical remodeling.

### Coupling surface mechanosensation to cell polarity

An important set of findings in this study is to discover that cells completely rewire polarity upon local CW damage. First, cells depolarize by detaching most polarity regulators from growing tips upon laser damage, which occurs even though the wound is formed at distance from cell tips. Second, they redirect polarized-based trafficking to the wound site to promote CW repair, and third, they repolarize at cell tips once the wound is repaired to reset polar growth. Some aspects of this polarity reorganization is consistent with previous similar laser-based assays in budding yeast [25], and was also documented in the context of cell compression [19]. Importantly, we found that local redirection of cell polarity to the wound is mediated by CW mechanosensors that reroute CW synthesis through Cdc42 and Rho1 based actin assembly and polarized trafficking. These findings suggests that CW mechanosensors might provide the upstream spatial cues required to redirect cell polarity, in addition to their role in promoting CW reinforcement trough the CIP. In addition to repair the CW, redirection of Cdc42-based polarity may also help promote local plasma membrane repair, by for instance localizing endocytosis and exocytosis to the wound site [26,36,59]. Our data suggest that mechanosensors signal directly or indirectly to Gef1 and Rgf1 to locally activate Cdc42 and Rho1. Interestingly, this occurs independently of classical MT or Tea1/Tea4 based pathways that promote *de novo* polarization to new ends at NETO (New End Take Off) [38,60,61] or to ectopic sites on cell sides in fission yeast [41,42,62]. The detailed mechanisms by which Wsc1 and Mtl2 sensors recruit these GEFs remain to be elucidated, but previous findings have documented a direct interaction between Wsc1 cytoplasmic C-terminal and the Rho1 GEF Rgf2 in fission yeast [10].

However, we found that the depolarization phenotype from cell tips was largely independent of mechanosensors and GEFs, as mutants in these factors still detached polarity zones while failing to recruit a new zone to wound sites (Fig 4, Fig 5 and Fig S5). In fission yeast, Sty1, a global stress activated MAPK, is thought to promote tip depolarization under various stress conditions, including osmotic and oxidative stresses, and could therefore also contribute to polar domain disassembly in this context [40,63,64]. Moreover, previous work conducted in S. *cerevisiae* showed that upon local CW damage Pkc1, an homologue of Pck1, triggers proteasomal degradation of polarity factors to resolve competitions between the bud and wound sites [25]. Finally, although dispensable for rerouting polarity to the wound site, we propose that the MT/Tea1 pathway may still be key to redirect polarity to incipient growing tips upon CW repair. Accordingly, we observed larger bulges alongside with long lasting residual GFP-Bgs4 signal at the wound under MBC treatment, as well as significant defects in polarity recovery to cell tips in *tea1*Δ mutant cells (Fig 4A and Fig S3C-D). Therefore, the assay implemented here may also help address some of the generic mechanisms regulating polarity competition and redirection.

### General relevance of CW repair mechanisms

In addition to enhance our understanding of mechanisms for local CW repair, we anticipate that our assay and findings may have broader relevance to other physiological situations. First, we speculate that CW repair as documented here, may have some level of similarity with CW closure during septation. In that view, the closing septum could be seen as a large wound that needs to be repaired and closed. Indeed, we documented a differential regulation of the two glucan synthase Bgs1 and Bgs4 by GEFs, reminiscent of that occurring during septum assembly [43,44]. In addition, we found that septating cells delay repolarization to the wound site until septation is completed (Fig S2H-J), suggesting that septum-based signals outcompete, laser-induced CW wound signals, much better than tip-based polarity. A possible explanation for this, is that local mechanical stresses or strains generated along the growing septa may be larger than at cell tips or wounds, retarding the rerouting of mechanosensors. Therefore, it would be interesting to understand if walled cells can use values of surface mechanical stress as a hierarchical system to prioritize the redirection of polarity and CW assembly [28].

Second, the connection we established between mechanosensation and polarity redirection during wound repair, may also have relevance to more natural polarity establishment, such as during NETO. Indeed, Wsc1 is generally abundant at new forming ends as a result of mechanical contacts between separating daughter cells, potentially serving as an early spatial cue for NETO [24]. In addition, *rgf*1Δ mutant cells are known to be monopolar, which based on our results, argues that this could in part be caused by decoupling surface mechanosensation with Rho1-based polarity [65]. Accordingly, a recent study suggests that Rgf1 has a dual influence on polarity, as it stabilizes Tea4 at cell tips but also acts independently of the Tea1/Tea4 module to prevent branching in *tea1*Δ mutant by stabilizing polarity at cell tips [66].

In other ecological contexts, the CW may be exposed to many insults from the environment such as compressive stress emerging from neighboring cells in dense biofilms or attacks from predacious yeasts or phytopathogenic fungi which can locally pierce the prey or host CW [7,67,68]. Further work documenting how these CW damaging events are being monitored, may broaden our appreciation for how surface mechanosensation, polarity and CW remodeling are orchestrated to ensure host cell survival during infection, or bypassed during predation to promote cell death.

## MATERIAL AND METHODS

### Fission yeast strains and growth conditions

Strains used in this study are listed in table S1. Standard *S. pombe* methods were used for yeast culture [69,70]. Cells were grown routinely on solid YE5S medium at room temperature (18-25°C) and switched at 25°C overnight in liquid YE5S before imaging. Cells growing in exponential phase were then transferred on 2 % agar YE5S agar pads for timelapse imaging. For shut-off experiments, cells were cultivated in solid EMM medium at room temperature and then switched at 25°C in liquid EMM supplemented with 5 µg/ml thiamine 16h before imaging [10]. Cells were then transferred on 2 % agar EMM agar pads supplemented with 5 µg/ml thiamine for timelapse imaging. Crosses were performed on ME medium or on nitrogen free EMM for thiamine sensitive strains. All EMM media were supplemented according to the strain needs. All crossed strains were segregated by tetrad dissection and selected on the relevant selecting media.

### Pharmacological inhibition

Chemicals used in this study are listed in table S2. Drug treatments were performed before timelapse imaging by modifying YE5S agar pad preparation with the corresponding drug concentrations. Cells were incubated on the modified agar pads 30 minutes at room temperature before imaging. Microtubule were depolymerized using 25 µg/ml Methyl 2-Benzimidazole Carbamate (MBC; Sigma-Aldrich) from a 100X stock dissolved in DMSO (Euromedex). F-actin was depolymerized using 100 µM Latrunculin A (LatA; Sigma-Aldrich) from a 100X stock dissolved in DMSO.

### Turgor pressure manipulation using sorbitol

Turgor pressure perturbations were performed by incubating cells 30 minutes before imaging on YE5S agar pads containing 0.5M sorbitol (Sigma-Aldrich). This sorbitol concentration is just above the isotonic concentration of 0.4 M described for *S. pombe*. Osmotic perturbations were all performed using *gpd*1Δ mutants, defective in turgor pressure recovery [47,48].

### Light microscopy

For all experiments, the temperature in the microscopy room was maintained at 23±1°C. Live-cell imaging was performed using a 100X oil objective (Nikon CFI Plan Apo DM 100X/1.4 NA) on a Nikon Ti-Eclipse equipped with a motorized stage, perfect focus system, Yokogawa CSU-X1FW spinning head, iLaunch laser illumination system (Gataca Systems) and a Teledyne Photometrics sCMOS Prime BSI camera. This confocal microscope was controlled using the MetaMorph software.

For live Cell Wall (CW) thickness measurement [9] cells expressing the plasma membrane marker GFP-LactC2 [71] were deposited on YE5S agar pads containing 10 µg/ml of lectins from *Griffonia simplicifolia* coupled with Alexafluor647 (Bs-IB_4_-Alexafluor647; ThermoFisher) to label the outer contour of the CW. Cells were incubated at room temperature for 15 minutes before imaging. Confocal midplane images of isolated cells were then acquired for later analysis. Images of 200µm TetraSpeck^TM^ microsphere (ThermoFisher) were acquired across the field of view to correct for chromatic shift during CW thickness analysis [9].

For the plasma membrane permeability assay using Propidium Iodide (PI) (ThermoFisher), cells were incubated 30 min before imaging at room temperature on a YE5S agar pad containing 10 µg/ml of PI.

### Laser irradiation

To perform laser irradiation, we used the iLas Pulse system in “targeted” mode with a 355 nm pulse laser (Gataca Systems) coupled to the confocal microscope described above and a 60X oil immersion objective compatible with UV-A (Nikon CFI Apo λS 60X Oil, 1.4 NA). Laser calibration was performed on a blue fluorescent slide (Chroma). Cells were irradiated on circular ROIs of a diameter of 1.8 µm centered on the cell periphery. The “8 repeats” setting was used for all laser irradiation experiments. Laser power and/or yeast cell sensitivity was variable across experiments. To wound the cells in a consistent manner, laser power sensitivity was tested on cells before irradiation experiments by increasing and decreasing the laser power by increments of 5%. The laser power used for experiments was the one causing no cell death within 5 minutes after irradiation, just below the laser power inducing short term cell death (see Fig 1). Typical laser power used on WT and single mutants was 50±10%. Laser power used on the *gef1*Δ *rgf1*Δ double mutant and the *wsc1*Δ *nmt1-mtl2* mutant in presence of thiamine was 35±5%. After laser irradiation, the objective was switched back to 100X for live imaging.

### Correlative Light Electron Microscopy (CLEM)

For CLEM imaging, cells were deposited on plasma-cleaner treated glass bottom petri dish with a numerated grid for image position registration (Mattek 35 mm Dish, No. 1.5 Gridded Coverslip), coated with 1 mg/ml poly-lysine (Sigma-Aldrich) and 0.1 mg/ml unlabeled lectin from *Griffonia simplicifolia* (Sigma-Aldrich). Cell were incubated 10 min at room temperature and rinsed twice with YE5S. Under the microscope, after choosing a position on the grid (Fig S1C), cells were ablated as described above. Just after laser irradiation a bright field image of the cells was taken using the 60X magnification and laser ROI positions were saved. 5 min after irradiation, cells were fixed with 1% glutaraldehyde in YE5S for 2h at 4°C. Samples were post fixed with 2% osmium tetroxide in water and progressively dehydrated using increasing ethanol concentration from 30% ethanol in water to 100% ethanol. Samples were then embedded in epoxy resin. 70 nm sections of the position of interest were generated using an ultramicrotome (Leica Ultracut UC6) and deposited on formvar/carbon coated copper grids. Sections were post-stained with 4% uranyl acetate and lead citrate in water. Samples were imaged on a FEI Tecnai12 transmission electron microscope at 120 kv equipped with a 4Kx4K OneView camera (Gatan). Partial deformations were often observed on cells, probably due to resin embedding, but in most cases the CW remained intact.

### Image analysis

Fluorescence intensities at the wound site and at the old end were measured in Fiji using rectangular ROIs of 14 x 50 pixels (0.75 x 2.69 µm) positioned at the cell surface. Built-in tools in Fiji were then used to subtract the background, and correct for photobleaching using the simple ratio method [72]. “Normalized fluorescence intensity”, as used throughout the manuscript, corresponds to the ratio between the measured fluorescence intensity at a given timepoint and the initial fluorescence intensity 5 min before laser irradiation. The “maximum enrichment” value corresponds to the mean of the maximum normalized fluorescence intensity obtained for each cell. “Time of maximum enrichment” corresponds to the mean of the time at which the maximum enrichment was obtained for each cell. For the cell permeability assay, PI fluorescence intensities were measured in Fiji using circular ROIs positioned on cell nucleus. “Normalized PI fluorescence” corresponds to the ratio between the measured fluorescence intensity at a given timepoint and the initial fluorescence intensity in the nucleus 5 min before laser irradiation.

Curvature measurements were done on images of cells expressing the GFP-LactC2 membrane marker. The Kappa plugin was used in Fiji to extract plasma membrane curvature at the wound site in WT and at the septum in the *ace2*Δ mutant by drawing 6 points along the plasma membrane. In the *ace2*Δ mutant, only the outer septa were analyzed because the inner septum tended to split. Cell length measurements were performed by measuring the long axis of the cell using phase contrast images. Growth rates were computed as the slope of cell length values plotted as a function of time within given time intervals.

To measure CW thickness after laser irradiation on Transmission Electron Microscopy (TEM) images, cells were first attributed to cells on bright field images using their position on the CLEM grid. Laser irradiation ROIs were then used to precisely determine the position of the wound site. CW thickness was measured in Fiji at the wound site and at the opposite side along a line perpendicular to the wound site and the long axis of the cell.

Live CW thickness measurements were carried out using a Matlab analysis pipeline described in a previous study [9]. The local distance between gaussian maximums of the far-red signal form lectins and the green signal form the membrane marker was computed along the cell contour to generate a CW thickness map. The chromatic shift between the two signals was corrected using a vectorial shift map of the field of view obtain form 200 nm TetraSpeck^TM^ microsphere images. Mean values of thickness along the wound site and along the corresponding CW length at the opposite site were extracted from the CW thickness map using the Matlab program. Mean values of thickness were used to compute the thickness ratio between the wound site and the opposite side. For CW thickness kinetics showing the evolution of this ratio after laser irradiation, the ratio was normalized on the initial ratio observed 5 min before irradiation.

For image display, brightness and contrast were adjusted in Fiji: the same parameters were used on images from the same timelapse. Excel (Microsoft) was used to calculate normalized values.

GraphPad Prism (GraphPad Software) was used to generate graphics and conduct the statistical analyses reported in figure legends.

## ACKNOWLEDGMENTS

We thank Celia Municio-Diaz and all members of the Minc laboratory for technical assistance and discussion. We acknowledge the ImagoSeine core facility of the Institut Jacques Monod, member of the France BioImaging infrastructure (ANR-24-INBS-0005 FBI BIOGEN) and GIS-IBiSA. This work was supported by the Centre National de la Recherche Scientifique (CNRS), the Université Paris Cité, and grants from La Ligue Contre le Cancer (EL2021.LNCC/ NiM), the Agence Nationale pour la Recherche (ANR, “CellWallSense” no. ANR-20-CE13-0003-02), and the Fondation Bettencourt Schueller (“Impulscience”), to N.M.

## AUTHOR CONTRIBUTIONS

Conceptualization, N.M., and Y.R.; Methodology, N.M., R. L.B., D. A.C., Y.R., Writing –Original Draft, N.M., and Y.R Draft Editing. N.M., R. L.B., Y.R.

