## Supplementary Material for "Mechanosensation Promotes Local Cell Wall Repair"

This supplementary file contains 5 supplemental figure and figure legends, 2 supplemental tables and 5 supplemental movie legends

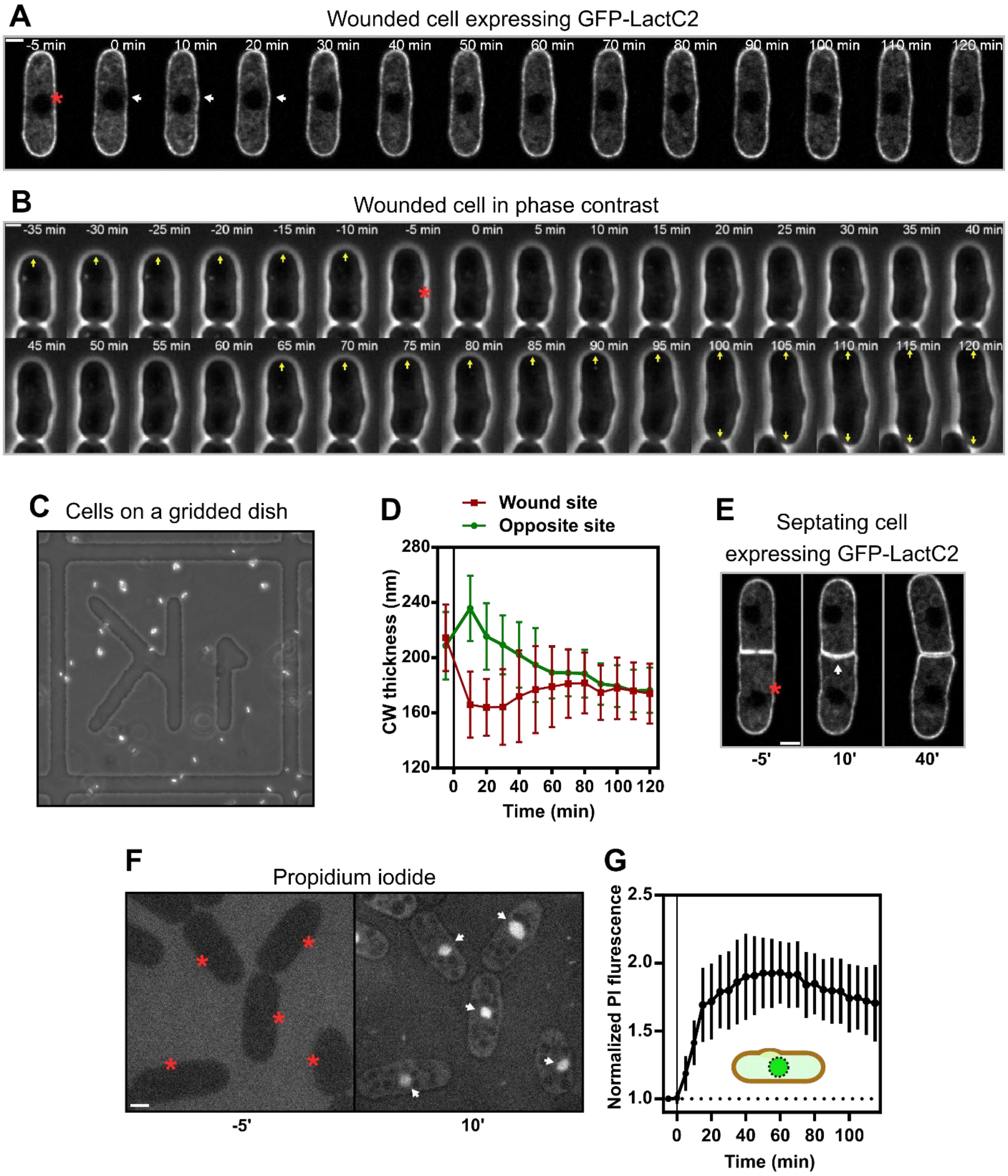

**Figure S1. Impacts of local cell wall damages created by laser irradiation**. **(A)** Confocal time-lapse of a cell expressing GFP-LactC2 before and after laser damage. Time zero was acquired few seconds after irradiation. The site of laser irradiation is marked by a red asterisk. White arrows point to the emerging bulge at the wound site. **(B)** Confocal time-lapse of a WT cell before and after laser damage. Yellow arrows point to growing cell tips. **(C)** Bright field image taken at low magnification of a gridded dish used for Correlative Live Electron Microscopy (CLEM). Yeast cells are visible as bright dots. **(D)** Kinetic of CW thickness at the wound site and at the opposite site extracted from CW thickness maps (n=24 cells). **(E)** Confocal timelapse of a septating cell expressing GFP-LactC2. The white arrow points to the septum that deforms towards the wounded cell. **(F)** Confocal images of cells labelled with 10 µg/ml Propidium Iodine (PI) before and after laser induced damage. White arrows point to PI-stained nuclei. **(G)** Kinetic of nuclear PI signal normalized to initial signal before irradiation in wounded cells (n=15 cells). Error bars represent SDs. Scale bars: 2 µm.

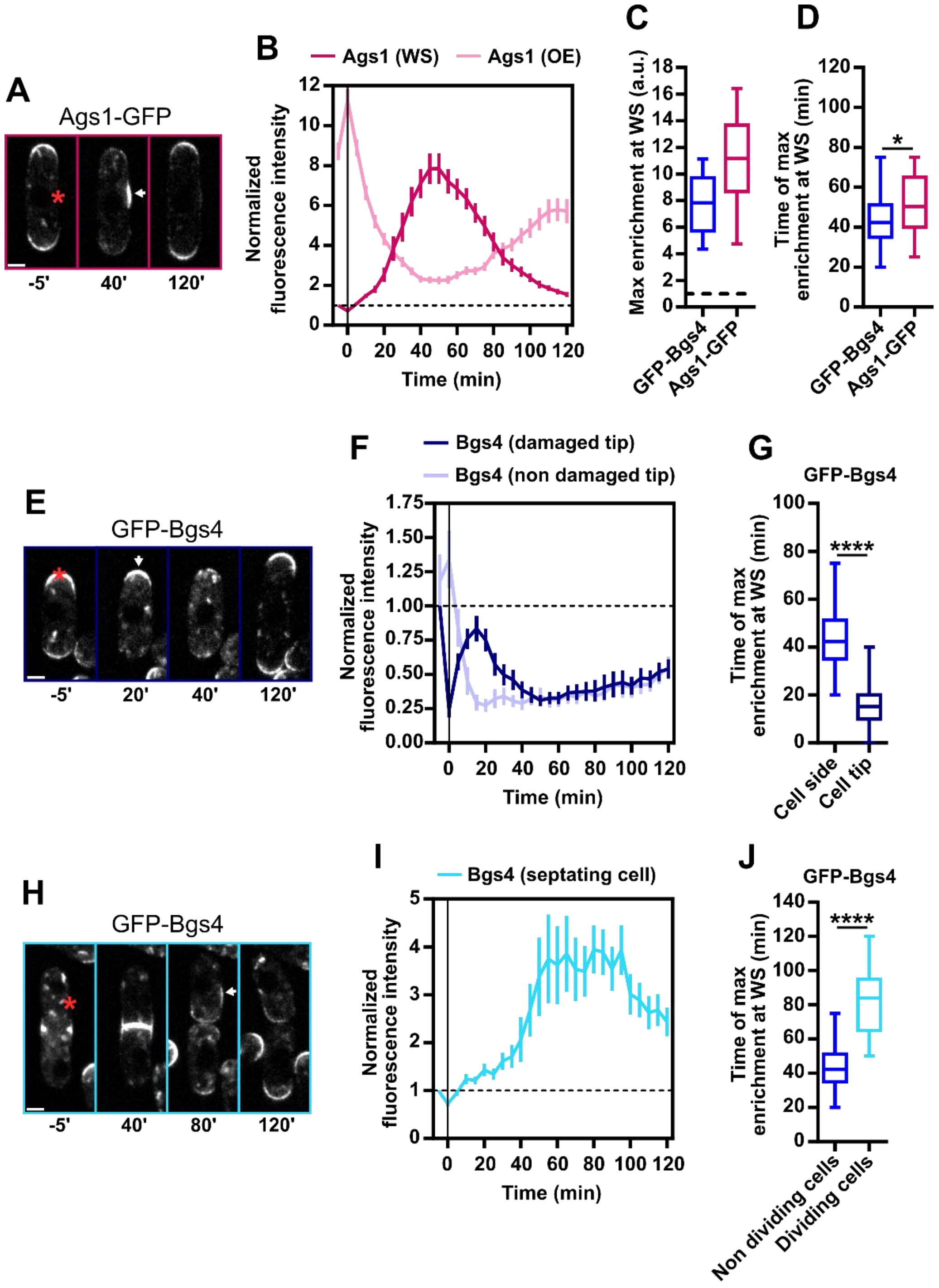
A
Ags1-GFP
-5'
40'
120'
B
Ags1 (WS)
Ags1 (OE)
Normalized fluorescence intensity
Time (min)
C
Max enrichment at WS (a.u.)
GFP-Bgs4
Ags1-GFP
D
Time of max enrichment at WS (min)
*
GFP-Bgs4
Ags1-GFP
E
GFP-Bgs4
-5'
20'
40'
120'
F
Bgs4 (damaged tip)
Bgs4 (non damaged tip)
Normalized fluorescence intensity
Time (min)
G
GFP-Bgs4
Time of max enrichment at WS (min)
****
Cell side
Cell tip
H
GFP-Bgs4
-5'
40'
80'
120'
I
Bgs4 (septating cell)
Normalized fluorescence intensity
Time (min)
J
GFP-Bgs4
Time of max enrichment at WS (min)
****
Non dividing cells
Dividing cells

**Figure S2. Glucan synthase recruitment to the wound site in different conditions. (A)** Confocal timelapse of a WT cell expressing Ags1-GFP before and after laser irradiation. Laser damage sites are marked by red asterisks. White arrows point to protein accumulation zones at the wound site (WS). **(B)** Kinetics of Ags1-GFP signal intensity at the WS and the old end (OE): time zero was acquired few seconds after laser irradiation (n= 30 cells). **(C)** Mean maximum enrichment of GFP-Bgs4 and Ags1-GFP measured at the WS. (n=30 cells for both strains). 1 correspond to the initial fluorescence signal measured at the WS before laser damage. **(D)** Mean timing of maximum GFP-Bgs4 and Ags1-GFP enrichment at the WS (n=30 cells). **(E)** Confocal timelapse of a cell expressing GFP-Bgs4 before and after laser irradiation at the old cell end. **(F)** Kinetics of GFP-Bgs4 signal intensity at the wounded and non-wounded cell tip (n=30 cells). **(G)** Mean timing of maximum GFP-Bgs4 enrichment at cell sides vs cell tips (n=30 and 27 cells respectively). **(H)** Confocal timelapse of a septating cell expressing GFP-Bgs4 before and after laser irradiation. **(I)** Kinetic of GFP-Bgs4 signal intensity at the WS (n= 15 cells). **(J)** Mean timing of maximum GFP-Bgs4 enrichment at the WS in dividing vs non-dividing cells (n=15 cells for each condition). Error bars in B,F and I correspond to standard error of mean (SEM). Whiskers in C,D G and J correspond to full data range. Results were compared using the Mann-Whitney test (*$p < 0.05$; ****$p < 0.0001$). Scale bars: 2 μm.

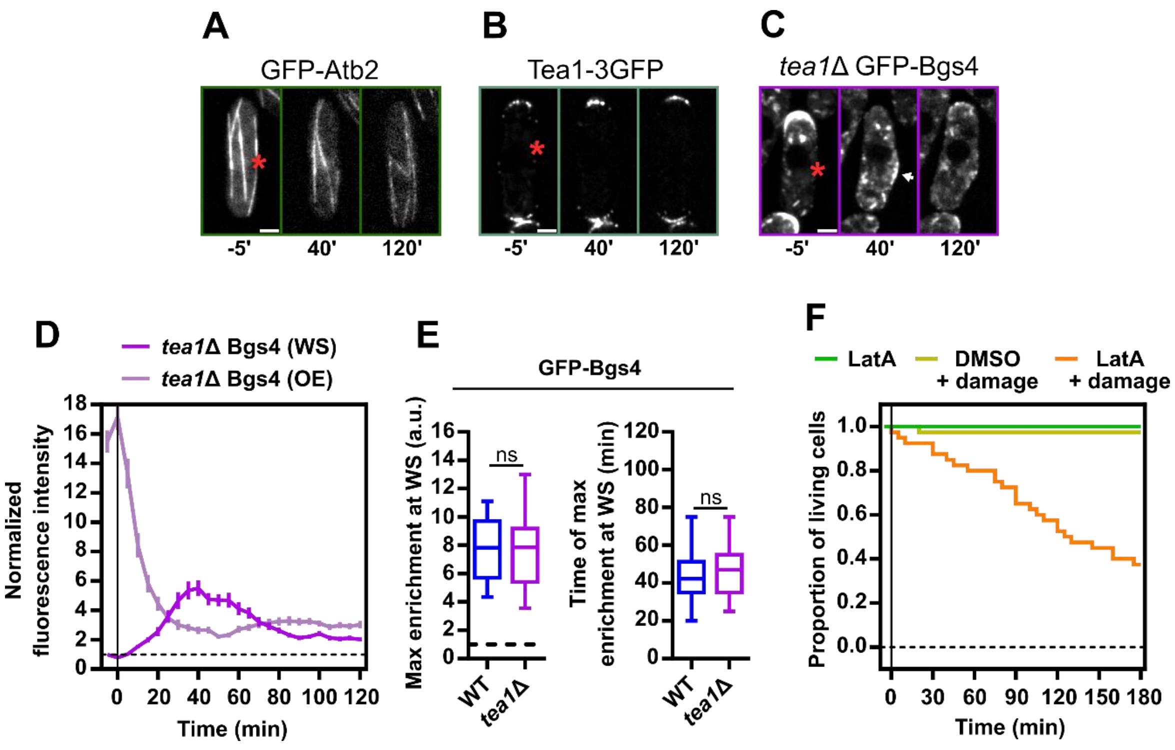
A
GFP-Atb2
-5'
40'
120'
B
Tea1-3GFP
-5'
40'
120'
C
tea1Δ GFP-Bgs4
-5'
40'
120'
D
tea1Δ Bgs4 (WS)
tea1Δ Bgs4 (OE)
Normalized fluorescence intensity
Time (min)
E
GFP-Bgs4
Max enrichment at WS (a.u.)
ns
WT
tea1Δ
Time of max enrichment at WS (min)
ns
WT
tea1Δ
F
LatA
DMSO + damage
LatA + damage
Proportion of living cells
Time (min)

**Figure S3. Microtubule and tea1-based polarity are dispensable for Glucan recruitment to wound sites. (A)** Timelapse of maximum intensity projections of a WT cell expressing GFP-Atb2 (to label microtubules) before and after laser irradiation. Laser damage sites are marked by red asterisks. **(B)** Confocal timelapse of a cell expressing Tea1-3GFP before and after laser irradiation. **(C)** Confocal timelapse of a *tea*1Δ mutant cell expressing GFP-Bgs4 before and after laser irradiation. The white arrow points to a protein accumulation zone at the wound site (WS). **(D)** Kinetics of GFP-Bgs4 signal intensity at the WS and the old end (OE) in *tea*1Δ mutant cells: time zero was acquired few seconds after laser irradiation (n= 28 cells). **(E)** Left panel: mean maximum enrichment of GFP-Bgs4 measured at the WS. Right panel: mean timing of maximum GFP-Bgs4 enrichment at the WS (n=30 and 28 cells). **(F)** Cell survival of: cells treated with 100 µM latrunculin A with no laser irradiation; cells treated with DMSO with laser irradiation; cells treated with 100 µM latrunculin A with laser irradiation. (n=40 cells for each condition). Error bars in D represent standard error of mean (SEM). Whiskers in E represent full data range. Results were compared using the Mann-Whitney test (ns: not significant). Scale bars: 2 µm.

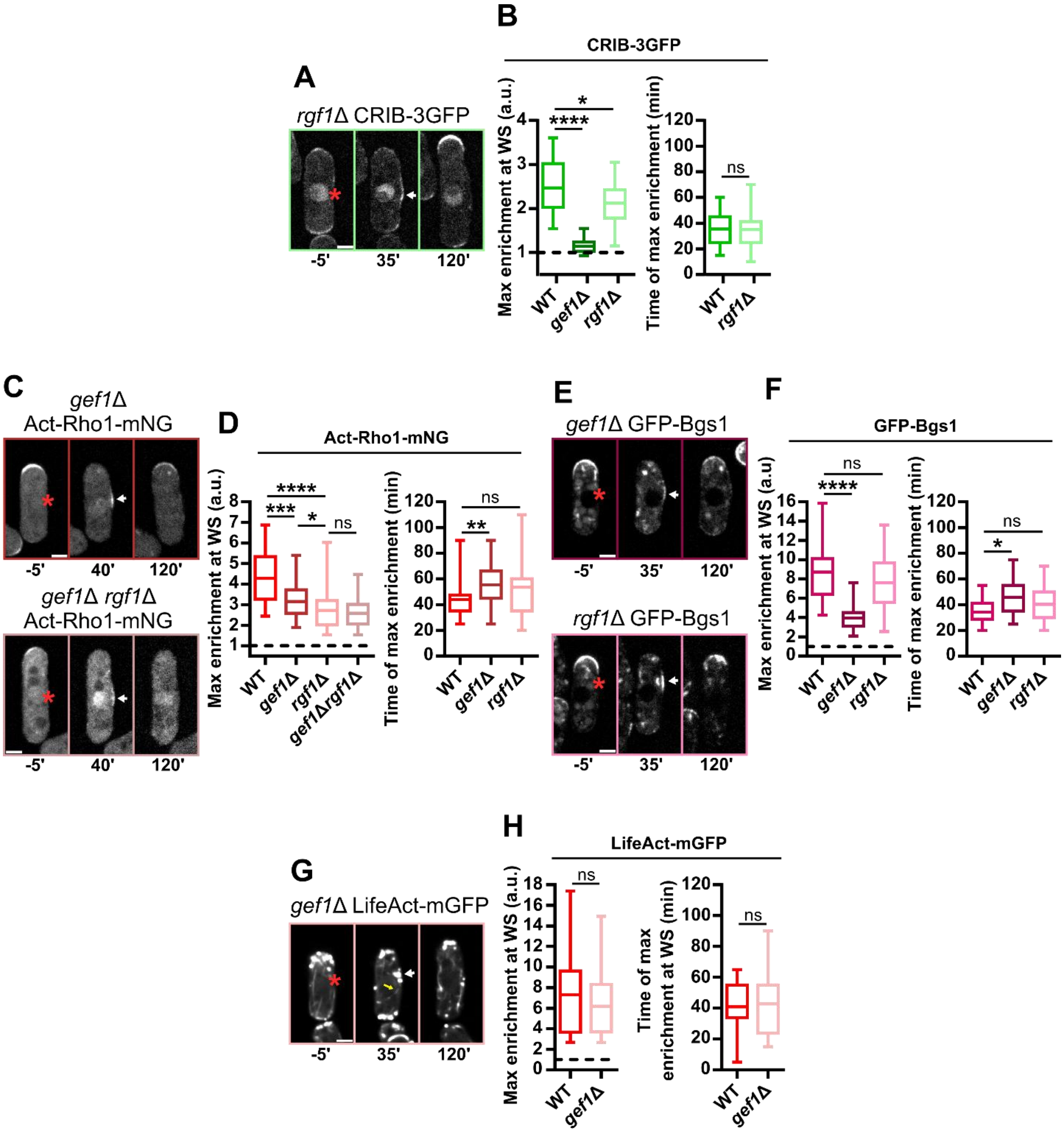

A
rgf1Δ CRIB-3GFP
-5'
35'
120'
B
CRIB-3GFP
Max enrichment at WS (a.u.)
Time of max enrichment (min)
WT
gef1Δ
rgf1Δ
****
*
ns
C
gef1Δ Act-Rho1-mNG
-5'
40'
120'
gef1Δ rgf1Δ Act-Rho1-mNG
D
Act-Rho1-mNG
***
gef1Δrgf1Δ
**
E
gef1Δ GFP-Bgs1
rgf1Δ GFP-Bgs1
F
GFP-Bgs1
Max enrichment at WS (a.u)
G
gef1Δ LifeAct-mGFP
H
LifeAct-mGFP
Time of max enrichment at WS (min)

**Figure S4. Gef1 and Rgf1 cooperate to regulate polarity and glucan synthases recruitment to wound sites. (A)** Confocal timelapse of a *rgf*1Δ mutant cell expressing CRIB-3GFP before and after laser irradiation. The white arrow points to a protein accumulation patch at the WS. Laser irradiation sites are marked by red asterisks. **(B)** Left panel: mean maximum enrichment of CRIB-3GFP measured at the WS. Right panel: mean timing of maximum CRIB-3GFP enrichment at the WS (n=30 cells in each strain). **(C)** Confocal timelapses of a *gef1*Δ mutant cell (top panel) and a *gef1*Δ *rgf1*Δ double mutant cell (bottom panel) expressing ActRho1-mNG before and after laser irradiation. **(D)** Left panel: mean maximum enrichment of ActRho1-mNG measured at the WS (n=30 cells in each strain). Right panel: mean timing of maximum ActRho1-mNG enrichment at the WS (n=30 cells in each strain). **(E)** Confocal timelapses of a *gef1*Δ mutant cell (top panel) and a *rgf1*Δ mutant cell (bottom panel), expressing GFP-Bgs1 before and after laser irradiation. **(F)** Left panel: mean maximum enrichment of GFP-Bgs1 measured at the WS. Right panel: mean timing of maximum GFP-Bgs1 enrichment at the WS (n=30 cells in each strain). **(G)** Confocal timelapse of a *gef*1Δ mutant cell expressing LifeAct-mGFP before and after laser irradiation. The yellow arrow points to an actin cable elongating from the WS. The white arrow points to actin patches at the wound site (WS). (**H)** Left panel: mean maximum enrichment of LifeAct-mGFP measured at the WS (n=30 cells). 1 correspond to the initial fluorescence signal measured at the WS before laser irradiation. Right panel: mean timing of maximum LifeAct-mGFP enrichment at the WS (n=30 cells). Whiskers represent full data range. Results were compared using the Mann-Whitney test (*p < 0.05; **p < 0.01; ***p < 0.001; ****p < 0.0001, ns: not significant). Scale bars: 2 µm.

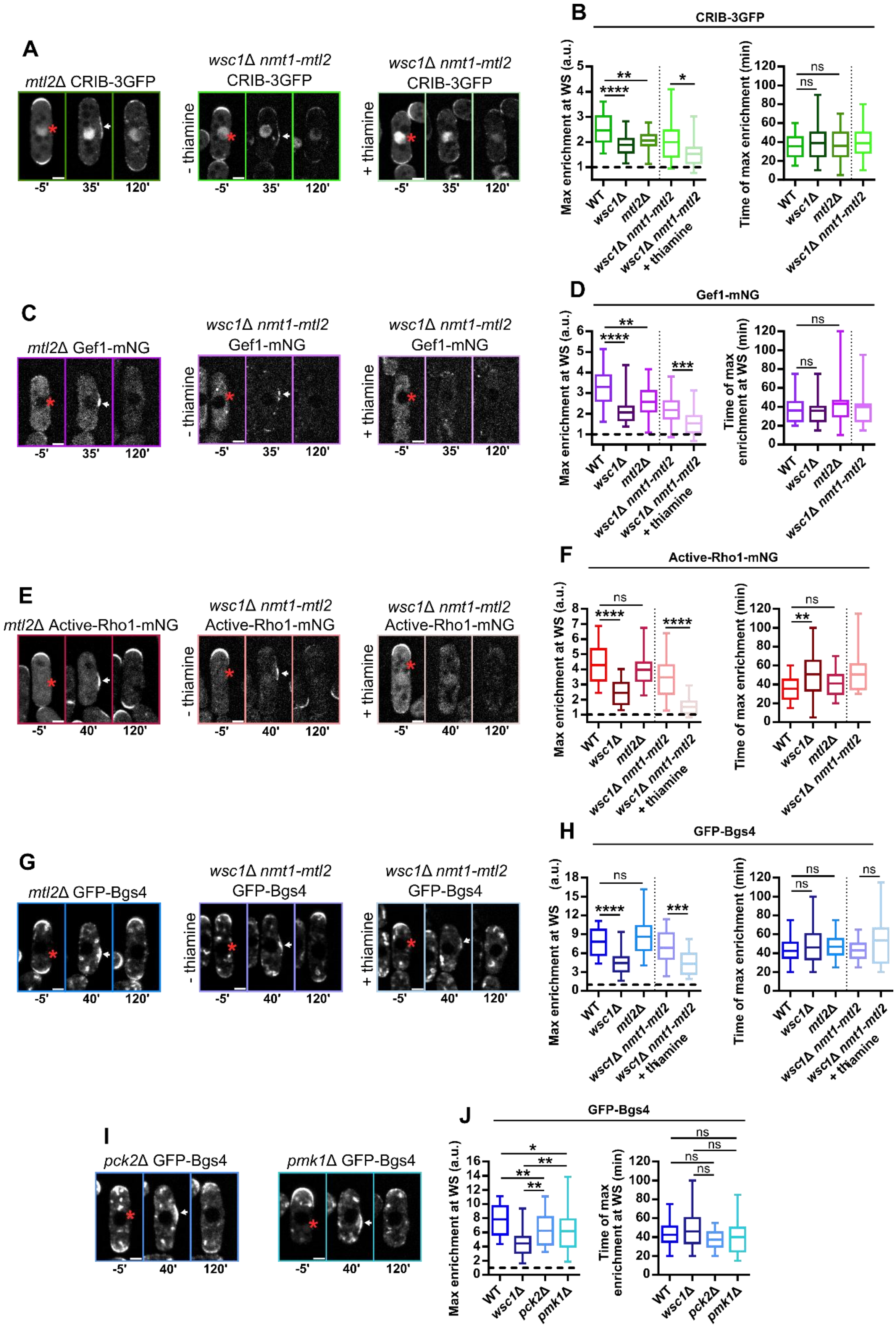

A
mtl2Δ CRIB-3GFP
wsc1Δ nmt1-mtl2
CRIB-3GFP
- thiamine
+ thiamine
-5'
35'
120'
B
CRIB-3GFP
Max enrichment at WS (a.u.)
Time of max enrichment (min)
WT
wsc1Δ
mtl2Δ
wsc1Δ nmt1-mtl2
wsc1Δ nmt1-mtl2 + thiamine
ns
C
mtl2Δ Gef1-mNG
wsc1Δ nmt1-mtl2
Gef1-mNG
D
Gef1-mNG
Time of max enrichment at WS (min)
E
mtl2Δ Active-Rho1-mNG
wsc1Δ nmt1-mtl2
Active-Rho1-mNG
-5'
40'
120'
F
Active-Rho1-mNG
G
mtl2Δ GFP-Bgs4
wsc1Δ nmt1-mtl2
GFP-Bgs4
H
GFP-Bgs4
I
pck2Δ GFP-Bgs4
pmk1Δ GFP-Bgs4
J
GFP-Bgs4
pck2Δ
pmk1Δ

**Figure S5. Mtl2 cooperate with Wsc1 to reroute polarity to the wound site independently of downstream SIP regulators.** **(A)** Confocal timelapses of a *mtl2*Δ mutant cell (Top panel); a non-induced *wsc1*Δ *nmt1-mtl2* mutant cell in EMM (Middle panel); a *wsc1*Δ *nmt1-mtl2* mutant cell induced with 5 µg/ml thiamine for 16h in EMM (Right panel), all expressing CRIB-3GFP. Laser irradiation sites are marked by red asterisks. The white arrows point to protein accumulation zones at the wound site (WS). **(B)** Left panel: Mean maximum enrichment of CRIB-3GFP measured at the WS. (n=33, 30, 30, 26, 27 cells, respectively). Right panel: mean timing of maximum CRIB-3GFP enrichment at the WS. (n=33, 30, 30, 26 cells, respectively). **(C)** Confocal timelapses of a *mtl2*Δ mutant cell (left panel); a non-induced *wsc1*Δ *nmt1-mtl2* mutant cell in EMM (middle panel); a *wsc1*Δ *nmt1-mtl2* mutant cell induced with 5 µg/ml thiamine for 16h in EMM (right panel), all expressing Gef1-mNG. **(D)** Left panel: mean maximum enrichment of Gef1-mNG measured at the WS. Right panel: mean timing of maximum Gef1-mNG enrichment at the WS (n=30 cells for all conditions). **(E)** Confocal timelapses of a *mtl2*Δ mutant cell (Left panel); a non-induced *wsc1*Δ *nmt1-mtl2* mutant cell in EMM (Middle panel); a *wsc1*Δ *nmt1-mtl2* mutant cell induced in 5 µg/ml thiamine for 16h in EMM (Right panel), all expressing ActRho1-mNG. **(F)** Left panel: mean maximum enrichment of ActRho1-mNG measured at the WS. Right panel: mean timing of maximum ActRho1-mNG enrichment at the WS (n=30 cells). **(G)** Confocal timelapses of a *mtl2*Δ mutant (left panel); an untreated *wsc1*Δ *nmt1-mtl2* mutant in EMM (middle panel); a *wsc1*Δ *nmt1-mtl2* mutant cell induced with 5 µg/ml thiamine for 16h in EMM (right panel), all expressing GFP-Bgs4. **(H)** Left panel: mean maximum enrichment of GFP-Bgs4 measured at the WS. Right panel: mean timing of maximum GFP-Bgs4 enrichment at the WS. (n=30 cells in all conditions). **(I)** Confocal timelapses of a *pck2*Δ mutant cell (left panel) and a *pmk1*Δ mutant cell (right panel), expressing GFP-Bgs4 before and after laser irradiation. **(J)** Left panel: mean maximum enrichment of GFP-Bgs4 measured at the WS (n=30, 30, 29, 28 cells respectively). Right panel: mean timing of maximum GFP-Bgs4 enrichment at the WS (n=30, 30, 29, 28 cells respectively). Whiskers represent full data range. Results were compared using the Mann-Whitney test (*p < 0.05; **p < 0.01; ***p < 0.001; ****p < 0.0001, ns: not significant). Scale bars: 2 µm.

| Strain | Source | Identifier |
|---|---|---|
| h+ leu1-32 ade6M210 ura4D18 | Minc lab stocks | AH146 |
| h90 shk1:GFP-lactC2:leu leu1-32 | Minc lab stocks | AH180 |
| h- ace2::kanMX shk1:GFP-lactC2:leu leu1-32 ura4D18 | This study | YR62 |
| h+ bgs1::ura GFP-bgs1:leu leu1-32 ura4D18 his3D1 | F. Chang lab stocks | FC1471 |
| h- bgs4::ura GFP-bgs4:leu leu1-32 ura4D18 his3D1 | F. Chang lab stocks | FC1472 |
| h- wsc1-mNG:kanMX leu1-32 ade6M210 ura4D18 | This study | AD1 |
| h- mtl2-mNG:kanMX leu1-32 ade6M210 ura4D18 | K L. Gould lab stocks | KGY19806 |
| h- rgf1::his3 leu:rgf1-GFP leu1-32 ade6M210 ura4D18 his3D1 | Y. Sanchez lab stocks | PG40 |
| h+ leu2:pck2:RBD-mNG:leu leu1-32 ade6M210 ura4D18 | M E. Das lab stocks | YMD1045 |
| h90 pck1::pck1-GFP-HA-kanR leu1-32 ade6-216 lys1-131 ura4D18 | NBRP yeast, Japan | FY14947 |
| h90 pck2::pck2-GFP-HA-kanR leu1-32 ade6-216 lys1-131 ura4D18 | NBRP yeast, Japan | FY15061 |
| h- CRIB-3GFP:ura leu1-32 ade6M210 ura4D18 | Minc lab stocks | NM123 |
| h- for3-3GFP:ura leu1-32 ade6M210 ura4D18 | Minc lab stocks | NM311 |
| h+ Pact1:LifeActin-mGFP:leu1 leu1-32 ade6M210 ura4D18 | M K. Balasubramanian lab stocks | MBY7519 |
| h90 myo52-3GFP::kanMX leu1-32 ade6M210 ura4D18 | Minc lab stocks | ST14 |
| h90 scd1-3GFP:kanMX scd2-mCherry:NatMX | Minc lab stocks | DB384 |
| h+ gef1-mNG:kanMX ade6M210 leu1-32 ura4D18 | M E. Das lab stocks | YMD910 |
| h- gef1::ura CRIB-3GPF:ura leu1-32 ade6M210 ura4D18 | This study | YR30 |
| h90 rgf1::kanMX CRIB-3GFP:ura ura4D18 | Minc lab stocks | AH306 |
| h+ gef1::ura leu2:pck2:RBD-mNG:leu leu1-32 ade6M210 ura4D18 | This study | YR44 |
| h+ rgf1::kanMX leu2:pck2:RBD-mNG:leu leu1-32 ade6M210 ura4D18 his3D1 | This study | YR35 |
| h- gef1::ura bgs1::ura GFP-bgs1:leu leu1-32 ade6M210 ura4D18 | This study | YR69 |
| h+ rgf1::kanMX bgs1::ura GFP-bgs1:leu leu1-32 ade6M210 ura4D18 his3D1 | This study | YR71 |
| h+ gef1::ura bgs4::ura GFP-bgs4:leu leu1-32 ade6M210 ura4D18 his3D1 | This study | YR33 |
| h- rgf1::kanMX bgs4::ura GFP-bgs4:leu leu1-32 ade6M210 ura4D18 | This study | YR15 |
| h+ rgf1::kanMX gef1::ura leu2:pck2:RBD-mNG:leu leu1-32 ade6M210 ura4D18 his3D1 | This study | YR56 |
| h+ rgf1::kanMX gef1::ura bgs4::ura GFP-bgs4:leu leu1-32 ade6M210 ura4D18 | This study | YR49 |

| | | |
|---|---|---|
| h- rgf1::kanMX gef1::ura shk1:GFP-lactC2:leu ade6M210 leu1-32 ura4D18 his3D1 | This study | YR60 |
| h- gpd1::kanMX wsc1-mNG leu1-32 ade6M210 | This study | YR58 |
| h+ gpd1::natMX CRIB-3GFP:ura leu1-32 ura4D18 | Minc lab stocks | AH312 |
| h+ gpd1::natMX leu2:pck2:RBD-mNG:leu leu1-32 ade6M210 ura4D18 | This study | YR83 |
| h- wsc1::kanMX CRIB-3GFP:ura ura4D18 | Minc lab stocks | AH291 |
| h90 mtl2::KanMX CRIB-3GFP:ura ura4D18 | Minc lab stocks | AH290 |
| h+ wsc1::ura kan-P81nmt1:mtl2 CRIB-3GFP:leu leu1-32 ade6M210 ura4D18 | This study | YR37 |
| h+ wsc1::kanMX leu2:pck2:RBD-mNG:leu leu1-32 ade6M210 ura4D18 his3D1 | This study | YR81 |
| h- mtl2::kanMX leu2:pck2:RBD-mNG:leu leu1-32 ade6M210 ura4D18 his3D1 | This study | YR79 |
| h- wsc1::ura kan-P81nmt1:mtl2 leu2:pck2:RBD-mNG:leu leu1-32 ade6M210 ura4D18 | This study | YR42 |
| h- gpd1::natMX bgs4::ura GFP-bgs4:leu leu1-32 | Minc lab stocks | VD181 |
| h- wsc1::kanMX bgs4::ura GFP-bgs4:leu leu1-32 | Minc lab stocks | VD179 |
| h- mtl2::his bgs4::ura GFP-bgs4:leu leu1-32 ade6M210 ura4D18 | This study | YR18 |
| h+ wsc1::ura kan-P81nmt1:mtl2 bgs4::ura GFP-bgs4:leu leu1-32 ura4D18 | This study | YR39 |
| h- ags1::ags1-GFP:leu:ura leu1-32 ade6M210 ura4D18 his3D1 | J C. Cotés lab stocks | #3166 |
| h- leu1-32:SV410:GFP-atb2:leu | Minc lab stocks | NM331 |
| h- tea1-3GFP:kanMX | Minc lab stocks | NM339 |
| h- rga4-GFP:kanR leu1-32 | P. Perez lab stocks | PPG1811 |
| h- tea1::natMX bgs4::ura GFP-bgs4:leu leu1-32 ade6M210 ura4D18 | Minc lab stocks | NM348 |
| h+ gef1::ura Pact1:LifeAct-mGFP:leu leu1-32 ade6M210 ura4D18 | This study | YR46 |
| h- ars1(BlpI):Padh13:CRIB-3mCitrine:leu2 leu1-32 ade6M210 ura4D18 | K. Sawin lab stocks | KS7311 |
| h- sty1::natMX bgs4::ura GFP-bgs4:leu leu1-32 ura4D18 | Minc lab stocks | AH440 |
| h+ wsc1::kanMX gef1-mNG:kanMX leu1-32 ade6M210 ura4D18 his3D1 | This study | YR67 |
| h+ mtl2::kanMX gef1-mNG:kanMX leu1-32 ade6M210 ura4D18 his3D1 | This study | YR65 |
| h- wsc1::ura kan-P81nmt1:mtl2 gef1-mNG:kanMX leu1-32 ade6M210 ura4D18 | This study | YR73 |
| h+ pck2::kanMX bgs4::ura GFP-bgs4:leu leu1-32 ura4D18 | This study | YR23 |
| h+ pmk1::KanR bgs4::ura GFP-bgs4:leu leu1-32 ade6M210 ura4D18 his3D1 | This study | YR17 |

### Tables S1. Fission yeast strains used in this study

| Chemical | Source | Identifier |
|---|---|---|
| Dimethyl sulfoxide (DMSO) | Euromedex | Cat# SC-358801 |
| Methyl 2-Benzimidazole Carbamate (MBC) | Sigma-Aldrich | Cat# 378674 |
| Latrunculin A (LatA) | Sigma-Aldrich | Cat# L5162 |
| D-Sorbitol | Sigma-Aldrich | Cat# S1876 |
| Thiamine hydrochloride | Sigma-Aldrich | Cat# T4625 |
| Conjugated lectin: Bs-$IB_4$-Alexafluor647 | Thermo Fisher | Cat# I32450 |
| Propidium iodide (PI) | Thermo Fisher | Cat# P3566 |

### Table S2. Chemicals used in this study

**Supplemental Movie Legend**

**Movie S1. CW thickness dynamic of a wounded cell.** Representative timelapse of CW thickness of a wounded WT cell expressing GFP-LactC2 and labeled with lectin coupled to Alexafluor647 (see methods). Laser irradiation sites are marked by red asterisks. The black arrow marks local CW thinning after laser irradiation. Laser irradiation was performed at t=0.

**Movie S2. Rerouting of CIP components and glucan synthases to the wound site.** Representative confocal timelapses of cells expressing, from left to the right: Wsc1-mNeonGreen, Rgf1-GFP, ActRho1-mNeonGreen and GFP-Bgs4, before and after laser irradiation. Laser irradiation sites are marked by red asterisks. Images were acquired every 5 minutes. The image at time 0 was acquired few seconds after laser irradiation. Scale bars: 2 µm.

**Movie S3. Scd1 localization, local Gef1 recruitment and Cdc42 activation after local CW damage.** Representative confocal timelapses of cells expressing, from left to the right: Gef1-mNeonGreen, Scd1-3GFP and CRIB-3GFP, before and after laser irradiation. Laser irradiation sites are marked by red asterisks. Images were acquired every 5 minutes. The image at time 0 was acquired few seconds after laser irradiation. Scale bars: 2 µm.

**Movie S4. Alteration of Bgs4 recruitment to the wound site in GEFs mutants.** Representative confocal timelapses of GEFs mutants expressing GFP-Bgs4, from left to the right: *gef1*Δ, *rgf1*Δ and *gef1*Δ *rgf1*Δ cells, before and after laser irradiation. Laser irradiation sites are marked by red asterisks. Images were acquired every 5 minutes. The image at time 0 was acquired few seconds after laser irradiation. Scale bars: 2 µm.

**Movie S5. Defects in Gef1 recruitment in mechanosensor mutants.** Representative confocal timelapses of mechanosensors mutants expressing Gef1-mNG: from left to the right: *wsc1*Δ, *mtl2*Δ and the *wsc1*Δ *nmt1-mtl2* shut-off in the absence or presence of thiamine (16h), before and after laser irradiation. Laser irradiation sites are marked by red asterisks. Images were acquired every 5 minutes. The image at time 0 was acquired few seconds after laser irradiation. Scale bars: 2 µm.

**Movie S6. Impaired Bgs4 recruitment to the wound site in mechanosensors mutants.** Representative confocal timelapses of mechanosensors mutants expressing GFP-Bgs4, from left to right: *wsc1*Δ, *mtl2*Δ and *wsc1*Δ *nmt1-mtl2* shut-off in the absence or in presence of thiamine (16h), before and after laser irradiation. Laser irradiation sites are marked by red asterisks. Images were acquired every 5 minutes. The image at time 0 was acquired few seconds after laser irradiation. Scale bars: 2 μm.
